# Zinc Sulfide-Based Hybrid Exosome-Coated Nanoplatform for Targeted Treatment of Glioblastoma in an Orthotopic Mouse Glioblastoma Model

**DOI:** 10.1101/2020.07.28.226076

**Authors:** Jingxin Mo, Meiying Li, Edith K.Y. Tang, Shaoyuan Li, Wei Zhao, J. Justin Gooding

## Abstract

Core-hybrid shell hydroxychloroquine (HCQ) loaded zinc sulfide (ZnS) nanoparticles were synthesized, characterized and evaluated for the treatment of glioblastoma cells *in vitro* and *in vivo*. These particles, denoted as HCQ@ZnS@exo@iRGD, consist of hollow ZnS nanoparticles loading with the autophagic inhibitor of hydroxychloroquine and covered by a hybrid shell containing exosomes (exo) and phosphatidylserine derived pH- and redox-responsive pegylated iRGD peptide, a gliomablastoma-homing and penetrating peptide. The hybrid exosomes enable HCQ@ZnS with good permeability across the blood-brain barrier and targeting ability to glioblastoma cells in orthotopic mouse glioblastoma model. ZnS acts as a photosensitizer for reactive oxygen species (ROS) production to inflict damage to organelles within glioblastoma cells. Hydroxychloroquine inhibits autophagic flux, which can subsequently lead to the accumulation of impaired organelles caused by the ROS. As a result, substantial selective damage to glioblastoma cells was realized owing to the hybrid exosomes guiding the anti-tumour effects of hydroxychloroquine and ZnS under light irradiation. The results provide evidence for the utility of HCQ@ZnS@exo@iRGD as a therapeutic strategy for glioblastoma.

## 1. Introduction

Glioblastoma is the most common malignant brain tumor in adults. But there is still a lack of effective therapy. The median survival of patients with glioblastoma is just over 1 year.[1] This poor prognosis is due, in part, to the blood-brain barrier (BBB) and unfavourable tumor microenvironment (TME) prevent anti-cancer drugs from penetrating deep into the tumor. The blood-brain barrier (BBB) is a highly selective semipermeable border of endothelial cells that prevents solutes in the circulating blood from non-selectively crossing into the extracellular fluid of the central nervous system where glioblastoma reside. [2] The tumor microenvironment (TME) is the environment around a tumor, including the surrounding blood vessels, immune cells, fibroblasts, signaling molecules and the extracellular matrix (ECM). [3] As such physical barriers like BBB and TME represent major challenges for targeted therapy. [4]

Photodynamic therapy, driven by the induction of reactive oxygen species (ROS) from a photosensitizer to damage cellular organelles, holds considerable promise for cancer therapy. [5] However, the efficacy of photodynamic therapy is compromized when treating glioblastoma. One of the major challenges is elevated autophagic activity in tumor cells allows them to survive photodynamic therapy. [6] Thus, therapeutics to limit autophagy is emerging as a promising target for glioblastoma. Hydroxychloroquine is an FDA-approved drug for efficiently inhibiting autophagy functions, [7] and may enhance the anticancer effects of photodynamic therapy. A challenge with hydroxychloroquine though is the nonselective distribution of the drug *in vivo*. [8] It is particularly difficult for hydroxychloroquine, which is highly hydrophilic, to pass the BBB, circumvent harsh tumor microenvironment, and reach the glioblastoma cells to exert its pharmacological effect. [9] Thus, methods are requried to deliver hydroxychloroquine selectively to glioblastoma cells for acting with photosensitizer to achieve better treatment of glioblastoma.

Zinc sulfide (ZnS) is an atypical transition-metal sulfide with various morphologies. [10] Among them, hollow ZnS has received considerable attention based on its capacity for higher drug loading. [11] Recently, There is increasing interesting at ZnS in the field of photodynamic therapy due to its strong light-response activity originated from its large energy gap (~3.7 eV). Hollow ZnS nanoparticles have two bands to induce ROS; One is the valence band and another is the conduction band. The valence band (h^+^) reacts with H_2_O to generate OH^•^ radical. And the conduction band (e^-^) reacts with oxygen to initiate a reduction process to produce ^1^O_2_ radicals. [12]

Exosomes are 30-120 nm microvesicles secreted by a variety of cells through exocytosis.[13] Exosomes consisting of a lipid bilayer that is capable of shielding its contents from the immune system. Previous studies have demonstrated exosomes excreted from tumour cells cultured *in vitro* will specifically accumulate in areas where homogenous parental cells are located *in vivo*. [14] Owing to this so-called homing ability, we try to exploit exosome in drug delivery for specifically delivering hydroxychloroquine loaded ZnS nanoparticles to glioblastoma cells and reduce off target distribution.

The 3-amino-acid sequence ArgGlyAsp (RGD) is an αVβ_3_ integrin selective peptide. The internalizing RGD (iRGD) is a recently developed cyclic RGD peptide.[15] The iRGD peptide is widely acknowledged as an efficient cell membrane penetration peptide targeting to αvβ3 integrins and neuropilin-1 receptors, which overexpress in many tumor cells including glioblastoma. [16] Thus, we hypothesize functionalizing ZnS nanoparticles with iRGD will improve BBB crossing for better glioblastoma therapy.

Taking advantages of both exosomes and iRGD, the designed hybrid exosome will contain exosomes derived from human U87 glioblastoma spheroids and iRGD ligand with pH- and redox-responsive linkers. The aim of this hybrid exosome is to increase efficiency for crossing BBB and capability to target glioblastoma cells as well. And the pH- and redox-responsive linkers are vital for integrating forementioned double targetabilities with great adaptability, enabling HCQ@ZnS@exo@iRGD transformed within tumor microenvironment for untimately fusing with glioblastoma cells.

In this study, we synthesized core-shell hydroxychloroquine nanoparticles, consisting of core of hollow ZnS nanoparticles covered by shell of dual-stimuli responsive hybrid exosome containing exosome and phosphatidylserine with pH- and redox-responsive pegylated iRGD ligand. These particles, denoted as HCQ@ZnS@exo@iRGD, were formulated to enable glioblastoma cells targeting photodynamic/chemo therapy.

As shown in Figure 1, hydroxychloroquine molecules are incorporated into the hollow ZnS core before coating with exosome excreted by human U87 glioblastoma spheroids (defined as HCQ@ZnS@exo). Subsequently pH- and redox-responsive pegylated iRGD-modified liposomes are merged with HCQ@ZnS@exo to give HCQ@ZnS@exo@iRGD. Grafting of iRGD assists HCQ@ZnS@exo to cross the BBB and reach the glioblastoma. Then, owing to high GSH concentration in the acidic tumour microenvironment, the iRGD shell will be dissociated from the surface of the drug delivery vehicle. Subsequently the exposed exosome membrane of HCQ@ZnS@exo will specifically fuse with the glioblastoma cells. Based on this design, we hypothesize substantial damage to organelles of glioblastoma cells will be achieved by a visible light source triggering the ROS from hollow ZnS nanopartilces. The suppressed autophagy activity by hydroxychloroquine, may further agitate the cytotoxicity of ROS. Consequently, significant targeted glioblastoma cells elimination will be achieved in orthotopic mouse glioblastoma model.

**Figure 1.**
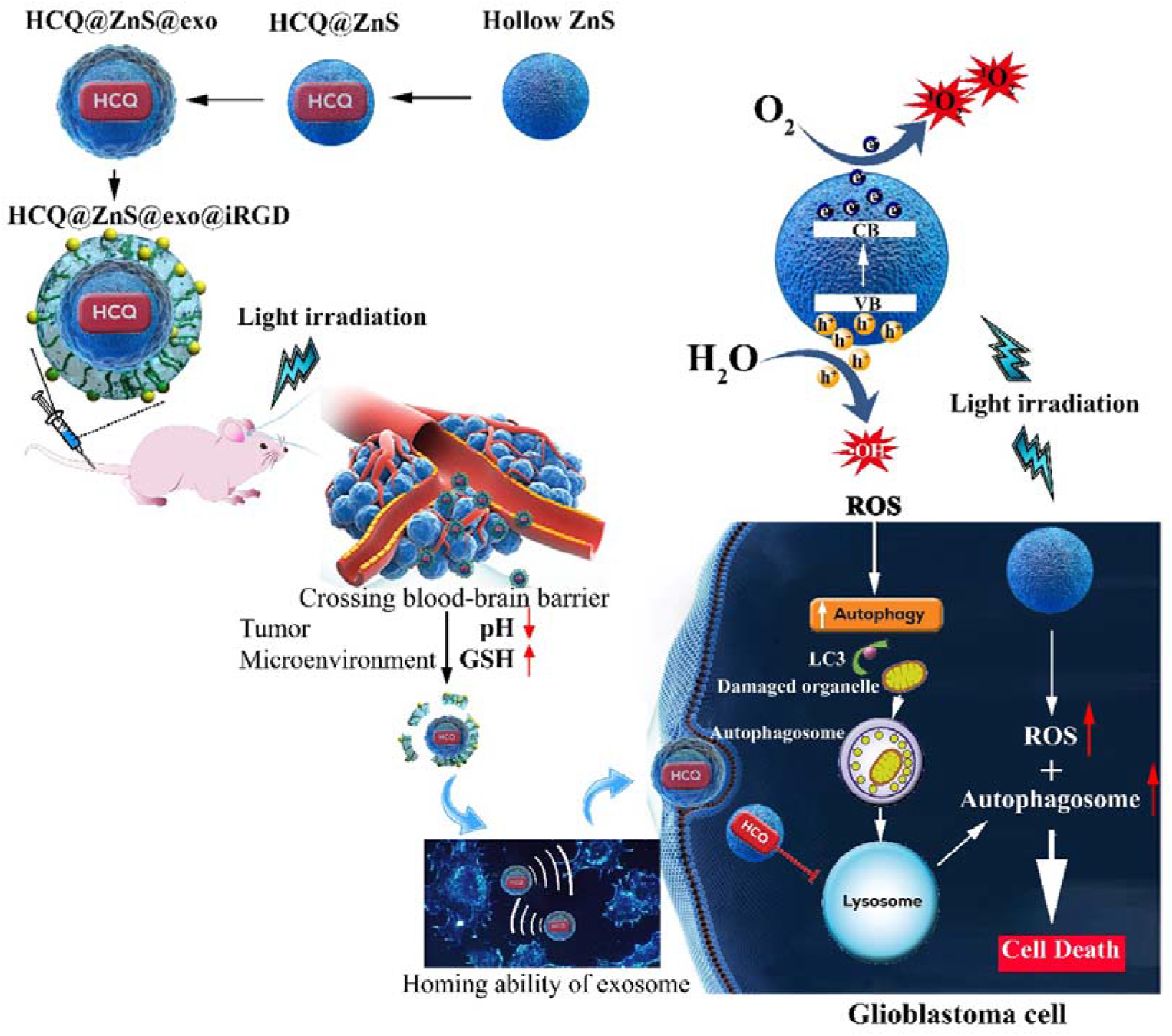
Schematic illustration of pH- and redox-responsive HCQ@ZnS@exo@iRGD as a photodynamic/chemo therapeutic nanoplatform for targeted treatment of glioblastoma cells in orthotopic mouse glioblastoma model. Abbreviations: ZnS: Zinc sulfide; HCQ: Hydroxychloroquine; Exo: Exosome; iRGD: phosphatidylserine with pH- and redox-responsive pegylated internalizing RGD ligand; GSH: Glutathione; CB: Conduction band; VB: Valence band; LC3: 1A/1B-light chain 3; ROS: Reactive oxygen species.

## 2. Methods

### 2.1. Materials

Zn (Ac)_2_·2H_2_O (AR) and gelatin (GR) were purchased from Aladdin Industrial Corporation (Shanghai, China). CH_4_N_2_S (AR) was purchased from Pesan Chemical Co.,Ltd (Changzhou, China). Phosphatidylserine and cholesterol were purchased from Eastar Chemical Corporation (Sacramento, USA). Hydroxychloroquine sulfate was obtained from MedChemExpress (NJ, USA). All the reagents were used without further purification.

### 2.2. Cell Culture

Human U87 glioblastoma cells were purchased from American Type Culture Collection (ATCC, Manassas, VA). Human MCF10A breast epithelial cells were obtained from Shanghai Cell Biology Institute of Chinese Academy of Sciences (Shanghai, China) and cultured in DMEM supplemented with 10% (v/v) FBS (ThermoFisher Scientific, Beijing, China) at 37°C in a humidified atmosphere of 5% CO2 after receiving. U87 cells in the phase of logarithmic growth were digested with trypsin (Gibco Biotech Co, Ltd, Beijing, China), collected, and resuspended in serum free DMEM. 10^6^ of U87 cells were seeded and cultured in 1O mL serum free stem cell culture media (EGF, 20 ng.mL; bFGF, 20 ng. mL; LIF, 10 ng.mL; B27, 1:50; PS, 1:100; Gibco) to form U87 cells sphere.[17] Culture supernatants were collected 2 days later and processed for exosome purification by differential centrifugation.[18]

### 2.3. Experimental animal

Female BALB/c nude mice weighing approximately 20 g were purchased from the Model Animal Research Center of Nanjing University (Nanjing, China). All animal experiments were performed in compliance with the Principles of Laboratory Animal Care (People’s Republic of China) and approved by the Ethics Committee of Guilin Medical University (ethics number GLMC-201803038).

### 2.4. Synthesis of ZnS hollow nanospheres

In a typical synthesis, Zn (OAc)_2_·2H_2_O (1 mmol) and CH_4_N_2_S (5 mmol) were dissolved in 30 mL distilled water (solution A). Gelatin (0.8780 g) was added to another 30 mL distilled water under vigorous magnetic stirring until the gelatin had dissolved (solution B). The two solutions were mixed and stirred for 10 min before being transferred to a Teflon-lined autoclave (100 mL in volume). The sample was heated at 130°C for 60 h. After cooling to room temperature naturally, the solids were collected by centrifugation, then washed with distilled water and absolute ethanol several times and dried at 60°C.[19]

### 2.5. ZnS characterization

X-ray powder diffraction (XRD) patterns were obtained on a Bruker D8 diffractometer using Cu Kα (λ = 1.5406 Å) radiation with an accelerating voltage of 40 kV and an applied current of 20 mA. The structure of the samples was analyzed by scanning electron microscopy (SEM: Hitachi S-4800) with an acceleration voltage of 5 kV and transmission electron microscopy (TEM: JEM-2100) with an acceleration voltage of 200 kV X-ray photoelectron spectroscopy (XPS) analysis was performed on a VG ESCALABMK II with a Mg Kα achromatic X-ray source (1253.6 eV).

### 2.6. *In vitro* methylene blue dye degraded with/without ZnS nanoparticles under different pH with/without visible light irradiation

Methylene blue (MB) dye degraded with/without ZnS was assayed using spectrophotometry as reported previously with modifications. [20] Briefly, 10 mg of ZnS was immersed in 250 mL of buffer solution containing MB (2mg/L) at different pHs (7.4, 5.5) with and without visible light irradiation from 600 W tungsten halogen lamp and stirred gently at 37°C. At each incubation time interval (up to 3 h), 500 μL of sample solution was collected and replaced with fresh buffer solution of the required pH value. The visible light-vis spectrum of the supernatant was recorded on a visible light-vis spectrophotometer (visible light-2450) to monitor the absorption behavior after centrifugation. The characteristic absorption peak of MB at 664 nm was used to determine the extent of its degradation.

### 2.7. Synthesis of phosphatidylserine with pH- and redox-responsive pegylated iRGD ligand

Phosphatidylserine with dual-stimuli responsive pegylated iRGD ligand (phosphatidylserine-*cis*-aconitic anhydride-cystamine-PEG-iRGD) was synthesized and characterized as detailed in *Supplementary information*. (Figure S1) Its chemical structure was confirmed by ^1^H-NMR (Mercury 400, Varian, Palo Alto) and FTIR (Spectrum One, Perkin Elmer, Waltham) shown in Figure S2.

### 2.8. Preparing HCQ@ZnS

Hydroxychloroquine was loaded into the hollow ZnS nanoparticles via physical encapsulation technique. Briefly, 10 mg of ZnS nanoparticles was immersed into a hydroxychloroquine phosphate buffer solution (pH 7.4) (5 mg/mL, 1 mL) and kept at 37°C for 24 h to achieve equivalent state. Subsequently, hydroxychloroquine loaded ZnS nanoparticles (HCQ@ZnS) were collected by centrifugation, washed with pure ethanol three times to remove the hydroxychloroquine on the outer surface of the ZnS nanoparticles. HCQ@ZnS was then freeze-dried for later experiments.

### 2.9. Synthesis of exosome coated HCQ@ZnS (HCQ@ZnS@exo)

Isolation/purification of exosome was based on differential centrifugation. [21] Specifically, 10 mL culture media was collected from U87 cells, cultured for 48 h in serum free stem cell culture media, and spun at 300 × g for 10 min. The supernatant was transferred to clean tube and spun at 2,000 × g for 15 min, filtered through a 0.8-μm filter before further ultracentrifuged at 100,000 × g for 90 min in an Optima L-90K, fixed angle Type 45 Ti rotor (k factor = 133; Beckman Coulter, Miami, FL, USA), at 4°C. After ultracentrifugation, pellets of samples were collected, and re-suspended in 1 mL PBS and used for next coating experiment. The protein content of exosome was quantified by BCA assay, using the manufacturer’s protocol.

HCQ@ZnS covered by exosome was synthesized via simple thin film hydration followed by membrane extrusion as described previously with minor modifications. [22] Briefly, 200 mg protein equivalent of purified exosome in 5 mL PBS was mixed with 20 mg HCQ@ZnS to form a thin film by rotavapor. It was then vortexed and sonicated for proper mixing after hydrated with PBS (pH 7.4). The mixture was extruded using 200 nm polycarbonate membrane filter to get nano sized HCQ@ZnS@exo, which were collected by centrifugation, washed three times with PBS (pH 7.4) and then freeze-dried for further use.

### 2.10. Synthesis of HCQ@ZnS@exo@iRGD

HCQ@ZnS loaded hybrid exosomes (HCQ@ZnS@exo@iRGD) were synthesized via simple thin film hydration followed by a membrane extrusion. Briefly, phosphatidylserine:phosphatidylserine-*cis*-aconitic anhydride-cystamine-PEG-iRGD (molar ratio of 5:1) and cholesterol in a molar ratio of 70:30 were dissolved with chloroform in a round-bottom flask attached to a rotavapor to form a thin film. Previously prepared HCQ@ZnS@exo in PBS were used to hydrate the dry lipid layer. Twenty mg protein equivalent of HCQ@ZnS@exo was added to 50 mg of lipid film in a final volume of 5 mL. It was then vortexed and sonicated (30% amplitude, 30 sec pulse on/off, for 2 min). The multilamellar HCQ@ZnS@exo@iRGD solution was extruded through a 200 nm polycarbonate membrane filter to get nano sized HCQ@ZnS@exo@iRGD, which were collected by centrifugation, washed three times with PBS (pH 7.4). Similarly, blank hybrid exosome was prepared as described in *Supplementary information*.

### 2.11. Characterization of hydroxychloroquine loaded ZnS nanoparticles

HCQ@ZnS, HCQ@ZnS@exo and HCQ@ZnS@exo@iRGD were characterized for particle sizes and surface charge via TEM and dynamic light scattering assay (DLS, Malvern ZSP). The surface morphology of the HCQ@ZnS, HCQ@ZnS@exo and HCQ@ZnS@exo@iRGD particles was characterized using scanning electron microscopy. Protein quantification of exosome and HCQ@ZnS@exo@iRGD were done by using Bradford assay (BioTek, Winooski, USA). The stability of the vesicles was studied by monitoring the change in size and PDI over a 7-day period.

### 2.12. Elucidation of fusion of lipid shell

Validation of fusion of HCQ@ZnS@exo@iRGD was performed *via* a Förster resonance energy transfer (FRET) study. Liposomes were prepared with FRET pairs: l-α-phosphatidylethanolamine-*N*-(4-nitrobenzo-2-oxa-1,3-diazole) (ammonium salt) (PE-NBD) (fluorescent donor, λem□=D525□nm) and l-α-phosphatidylethanolamine-*N*-(lissamine rhodamine-B sulfonyl) (ammonium salt) (PE-Rh-B) (fluorescent acceptor, λem□=□595░nm) (Merck KGaA, Darmstadt, Germany) in 1:7 DM ratio. FRET liposomes were synthesized by the thin-film method as described in previous research. [23] FRET fluorophore lipids, NBD acting as an electron donor and Rh-B acting as an electron acceptor, resulted in the formation of FRET liposome. For fusion analysis, 100 μL of HCQ@ZnS@exo@iRGD (1□mg/mL) was added to 20 μL FRET liposomes (EJmg/ml), mixed and bath sonicated for 5 min to initiate fusion. FRET liposomes, before and after fusion of hybrid exosome (HCQ@ZnS@exo@iRGD), were analyzed by fluorescence spectroscopy by exciting samples at 470 □nm and measuring the emission spectra between 500 and 700 □nm. Protein characterization of parental cells and exosomes and hybrid exosome (HCQ@ZnS@exo@iRGD) were done via SDS-PAGE analysis, dot blot assay, and Western blot.

For SDS-PAGE analysis, cell lysate, exosomes and hybrid exosome (HCQ@ZnS@exo@iRGD) were mixed with sample loading buffer (1:1) to give a final protein concentration of 200□μg/mL, respectively. The mixture was incubated at 90°C for 7 min, and 25 μL of each sample was loaded into the wells of 4-20% Mini-PROTEAN^®^ TGX Protein Gels. The gel was stained imaged by Bio-Rad imager (Kodak).

For dot blot assay, a drop of each sample (2-3 ≥ μL) was added to Polyvinylidene Fluoride (PVDF) membrane. The membrane was incubated with blocking buffer for 30 min at room temperature and then treated with primary antibodies CD81, CD63, CD11b, CD9, TSG101, β-actin, and histone H3 (Santa Cruz). After overnight incubation with the primary antibody, the membrane was washed with wash buffer and incubated with HRP conjugated anti-mouse IgG secondary antibody (Cell signaling). The membrane was further developed using Signal Fire ECL TM Reagent and immediately imaged by Bio-imager.

For Western blotting, an aliquot of 25 ? μL of sample mixed with sample loading buffer (1:1) containing 25□μg protein were loaded onto the gel. After completion of the SDS PAGE, the gel was transferred to PVDF membrane for the transfer of protein by the Western-blot method as described in our recent publications. [24] PVDF membrane was treated with primary antibodies β-Actin and CD9, CD63, hsp 70, LC3-I and II, SQSTM1/p62, GADPH (Santa Cruz) along with HRP conjugated anti-mouse IgG secondary antibody (Cell signaling). The membrane was further developed using Signal Fire ECL TM Reagent and immediately imaged for chemiluminescence by Bio-imager (Kodak).

### 2.13. *In vitro* study

#### 2.13.1. Cellular internalization of HCQ@ZnS@exo@iRGD

Cellular internalization of vesicles was studied using Confocal Laser Scanning Microscope (Carl Zeiss, LSM-700). 10,000-20,000 U87 cells and MCF10A cells were seeded in 8 well plates for 24 h at 37°C in 5% CO_2_ environment to form mono-layer cells. To form three-dimensional tumor spheroids, U87 cells were seeded at a density of 5×10^3^ cells per well in a Corning^®^ 96 Well Ultra Low Attachment Microplate. Certain amount of Rhodamine-B (Rh-B) instead of hydroxychloroquine for co-incubated with ZnS and then covered by hybrid exosome to prepare Rh-B labeled formulations at final concentration of 0.01 mg/mL. Cells were incubated with Rh-B labeled formulations for 3 h at 37°C with 5% CO_2_. Cells were then fixed by adding 4% (v/v) paraformaldehyde. The nuclei were stained with DAPI and cells were observed under a confocal laser scanning microscope. Image-Pro Plus 7 software (Media Cybernetics Inc., Rockville, MD, USA) was used to quantify the fluorescent intensity of internalized Rh-B labeled formulations.

#### 2.13.2. Cytotoxicity evaluation

Cytotoxicity of different formulations was analyzed on U87 cells using MTT assay. Briefly, 1×10^4^ cells were seeded in 96 well plates for 24□h at 37°C and 5% CO2. After that, cells were incubated with free hydroxychloroquine or different hydroxychloroquine formulations at varying concentration range (6.25, 12.5, 25, 50, 100, and 200 □μM) for 4Mh with/without visible light irradiation. Control cells were maintained with media only. MTT solution (20 μL, 5 mg/mL) was added to each well and cultures were incubated for another 4 h. DMSO was added to dissolve the insoluble formazan crystal formed after MTT treatment, and absorbance was recorded at 550 nm using microplate reader (BioTek, Synergy H1 Hybrid reader). IC_50_ values were determined from these absorbance measurements.

#### 2.13.3. Expression of GFP-LC3 in U87 cells

pBABE retroviral vector containing GFP-LC3 (Addgene, 22405) was transfected into PT67 cells (ATCC) using Lipofectamine 3000 (Life Technologies). [25] Retrovirus-containing suspension was collected after 48 □h, passed through a 45 μm filter to obtain viral particles and U87 were transduced in suspension while spinning at 180 *g* for 1 □h and subsequently selected with puromycin.

#### 2.13.4. Fluorescent imaging of lysosomes and autophagosomes

For visualization of lysosomes, U87 cells were treated for 3 h with free hydroxychloroquine (HCQ), HCQ@ZnS, HCQ@ZnS@exo or HCQ@ZnS@exo@iRGD at 40 μM hydroxychloroquine and labeled for 15 min with 25 nM LysoTracker Red (Invitrogen/Thermo Fisher Scientific, L7528). The control group of U87 was treated with PBS only. For visualization of autophagosomes, U87 cells expressing EGFP-LC3 were treated as above. The fluorescence signal was examined using confocal microscopy. To quantify de-acidification of lysosomes and autophagosome accumulation in cells, the area of lysosomal red puncta and the number of green (autophagosomes) puncta were quantified in at least 25 cells, respectively. Three independent experiments were performed. Lysosome and autophagosome were quantified using Image-Pro Plus 7 software.

### 2.14. *In vivo* study

#### 2.14.1. Establishment of orthotopic glioblastoma xenograft in BALB/c nude mice

The orthotopic mouse glioblastoma model were established through intracranial implantation of U87 cells as previously reported. [26] Briefly, 2 × 10^5^ U87 cells expressing luciferase (Luc-U87) were injected intracranially into the frontal lobes of BALB/c nude mice. The stereotactic coordinate of the injection spot on the mice was 0.5 cm posterior to the frontline, 0.5 cm to the left of the middle and 0.5 cm below the skull. Multiple holes were drilled around the injection site allowing for light penetration. The growth of orthotopic tumors were detected and quantified by bioluminescence imaging performed in an *In Vivo Image System* (IVIS) Spectrum (PerkinElmer, USA) by intraperitoneal injection of D-luciferin potassium salt (150 mg/kg).

#### 2.14.2. Different formulations’ distribution in glioblastoma bearing mice model

The mice bearing orthotopic glioblastomas were given free Cy 5.5, Cy 5.5@ZnS, Cy 5.5@ZnS@exo or Cy 5.5@ZnS@exo@iRGD via tail vein injection (*n*=3). The mice were then subjected to IVIS imaging system at various time points, revealing by the fluorescence of Cy 5.5. At 24 h post-injection, the mice were sacrificed. Major organs including brain were obtained for fluorescent imaging.

#### 2.14.3. Anti-glioblastoma, anti-stemness and pro-apoptosis effects *in vivo*

The mice bearing intracranial Luc-U87 glioblastoma were randomly divided into 5 groups (n = 9), and tail intravenously administered with free HCQ, HCQ@ZnS, HCQ@ZnS@exo or HCQ@ZnS@exo@iRGD with/without light irradiation and saline, respectively. The growth of the orthotopic glioblastoma was detected by the luciferin for *in vivo* imaging. (*n*=6) The injection treatment was carried out every 3 days for a total of eight injections. Six hours later after the injection, the tumor sites in the mice head were locally irradiated by 500 W tungsten halogen lamp for 10 min. Meanwhile, the body weights of the mice were determined every 3-4 days. The survival times were calculated since the glioblastoma inoculation to the day of death or until day 80, and the statistical difference was plotted by the Kaplan-Meier method.

Anti-stemness and pro-apoptosis effects, acute side effects of free HCQ, HCQ@ZnS, HCQ@ZnS@exo or HCQ@ZnS@exo@iRGD with/without light irradiation and saline on glioblastoma mice model was studied by immunofluorescence and biochemical analysis (*n*=3). On day 14, mice were euthanized. Tumors were cut into 5.0 μm thick slices, rinsed with PBS three times, incubated in binding buffer and stained with annexin V and propidium iodide (Lianke Technology, Hangzhou, China) for 15 min at 25°C under low-light conditions. Samples were washed again, stained with 5 mM DAPI. Tumor slices were also subjected to antigen retrieval in a pressurized heating chamber (Pascal, Dako, Denmark) in tris-EDTA (pH 9) at 115 °C for 1 min before incubation with primary antibodies against CD133 and CD90, which are stemness-related markers of glioblastoma. Slices are analyzed under a confocal microscope (Leica TCS SP5, Germany). Blood sample was collected from posterior vena cava. And an aliquot of each whole blood sample was centrifuged at 1,006 g for 10 min to obtain serum to determine the biochemical parameters. Hematological and biochemical analysis were conducted using a blood autoanalyzer (CDC Technologies, Ohio, USA).

### 2.15. Statistical analysis

Differences were assessed for significance using Student’s *t* test between two treatment groups or one-way analysis of variance among multiple groups. Three significance levels were reported: \**p* < 0.05, \*\**p* < 0.01 and \*\*\**p* < 0.001.

## 3. Result

### 3.1. Characterization of the ZnS nanoparticles

ZnS was synthesized using a hydrothermal reaction of Zn (OAc)_2_ and thiourea in the presence of gelatin at 130°C for 60 h (mass ratio of gelatin and Zn (OAc)_2_·2H_2_O is 4:1). Figure 2a shows the XRD pattern of the ZnS sample. Three peaks can be observed at 2*θ* = 28.8°, 48.4°, 57.4°, which can be indexed to (002), (110) and (112) diffraction peaks of the hexagonal zinc sulfide (No. 80-0007). No other impurity peaks were found. The morphology of the sample was observed by SEM and TEM analysis. From the SEM image (Figure 2b), we can see that the ZnS sample is composed of uniform nanospheres with a size of about 70-90 nm. The nanospheres are well dispersed with no obvious signs of aggregation. The low magnification TEM (Figure 2c) reveals the hollow spheres possess a rough surface. The amplified TEM image (Figure 2d) further demonstrates the hollow structure of ZnS. The high resolution TEM (HRTEM) image shows the lattice fringe spacings at about 0.31 nm, corresponding to the planar distance of the (002) plane of hexagonal ZnS (Figure 2d and inset).[27] XPS spectra of the ZnS sample was also measured. The XPS survey scan in Figure S3 shows the presence of Zn, S, N, C and O elements. The abundant N and C elements in ZnS should derive from surface-absorption species released by gelatin. All the binding energies indicate the formation of ZnS.

**Figure 2.**
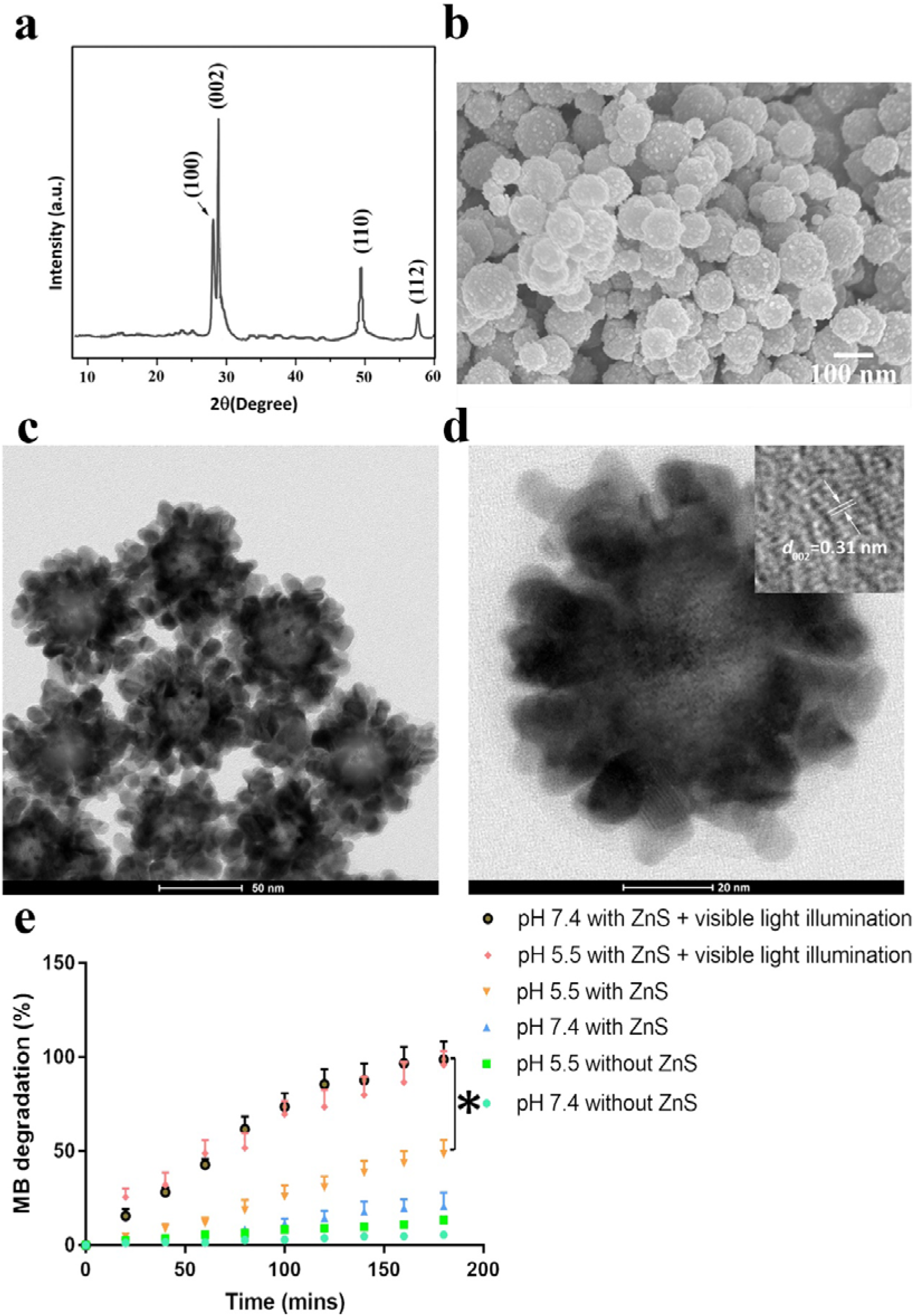
**(a)** XRD pattern, **(b)** SEM image, **(c)** TEM and **(d)** Amplified TEM images of ZnS. (inset: lattice fringes of ZnS (002)) **(e)** The profile of MB (methylene blue) degradation when co-incubated with the ZnS nanoparticles was investigated at different pH values with/without light illumination. Data shown are mean ± SD of 3 replicates. *, *p* < 0.05.

### 3.2. Photocatalytic activity of the ZnS nanoparticles

To confirm the photocatalytic performance of the ZnS nanoparticles we first looked at ability of the ZnS nanoparticles to degrade the dye methylene blue. The degradation profile of methylene blue with/without ZnS nanoparticles investigated at different pH values with/without light illumination are shown as Figure 2e. Under both pH 5.5 and pH 7.4 conditions, the percentage degradations of MB were above 95 % after 180 mins under light irradiation when containing ZnS nanoparticles. In comparison, under neutral conditions without light irradiation, the mean percentage degradation of methylene blue assisted by ZnS after 3 h was only 20.8%. Decreasing pH to 5.5 will increase MB degradation to 48.4% at the same scenario. Finally, in absence of ZnS nanoparticles, it is clearly seen that light illumination only caused less than 14% methylene blue conversion at pH 5.5 and 7.4. These results demonstrate that, under visible light illumination, the degradation of methylene blue significantly increased due to the high photocatalytic performances of ZnS.

### 3.3. Characterizations of HCQ@ZnS, HCQ@ZnS@exo and HCQ@ZnS@exo@iRGD

The chemical structure of phosphatidylserine-*cis*-aconitic anhydride-cystamine-PEG-iRGD and cholesterol, and graphic scheme of preparing blank hybrid exosome were shown in Figure S4. Small extracellular vesicles (sEVs) were harvested from conditioned media of U87 tumor spheroids as described in the method. TEM image showing a morphological characteristic of purified sEVs was presented as Figure S5a.

The products of HCQ@ZnS, HCQ@ZnS@exo and HCQ@ZnS@exo@iRGD were characterized for size, surface property, and protein content. A comprehensive result on the size distribution and zeta potential along with entrapment efficiency and drug loading of these nanoparticles is listed in Table 1. Particle sizes of HCQ@ZnS and HCQ@ZnS@exo was found to be 81□±□7□nm (PDI□=□0.17□±□0.04) (Figure 3a) and 99□±□9□nm (PDI□=□0.21 Ū±Ū0.02) (Figure 3b) with zeta potential of 2□±□0.4 and −15□±□1□mV, respectively. After modification of the HCQ@ZnS@exo with synthetic liposomes to give HCQ@ZnS@exo@iRGD these values changed to 116□±□18□nm (PDI□=□0.25□±□0.05) (Figure 3b) and −26□±□4□mV, respectively. The increase in the size of HCQ@ZnS@exo@iRGD is attributed to the insertion the synthetic liposomes into the bilayer of the exosomes which increased the interaction points of water molecule thereby increasing the hydration layers. Although the size of HCQ@ZnS@exo@iRGD is larger than HCQ@ZnS and HCQ@ZnS@exo, the most important factor is the homogeneity in the size distribution, which can be indexed by determining the polydispersity index (PDI). The PDIs of the three formulations are below 0.3, which indicate fairly good size homogeneity of these formulations.

**Table 1.** Physical characterizations of different ZnS based nanoparticles.

| Sample | Diameter<br>[nm] | PDI | Zeta potential<br>[mV]<sup>b</sup> | EE [%]<sup>c</sup> | DL [%]<sup>c</sup> |
| --- | --- | --- | --- | --- | --- |
| **ZnS** | 82±8<sup>a</sup> | 0.19±0.03<sup>a</sup> | -2.7±2<sup>b</sup> | N.A. | N.A. |
| **HCQ@ZnS** | 81±7<sup>a</sup> | 0.17±0.04<sup>a</sup> | 2±1<sup>b</sup> | 27.10±2.21 | 15.20±2.31 |
| **HCQ@ZnS@exo** | 99±9<sup>b</sup> | 0.21±0.02<sup>b</sup> | -15±1<sup>b</sup> | 97.52±5.6 | 6.45±1.85 |
| **HCQ@ZnS@exo@iRGD** | 116±18<sup>b</sup> | 0.25±0.05<sup>b</sup> | -26±4<sup>b</sup> | 94.66±7.9 | 5.05±1.34 |

DL: drug loading; EE: encapsulation efficiency; PDI: polydispersity index.
N.A. not applicable
n=5, mean±SD

**Figure 3.**
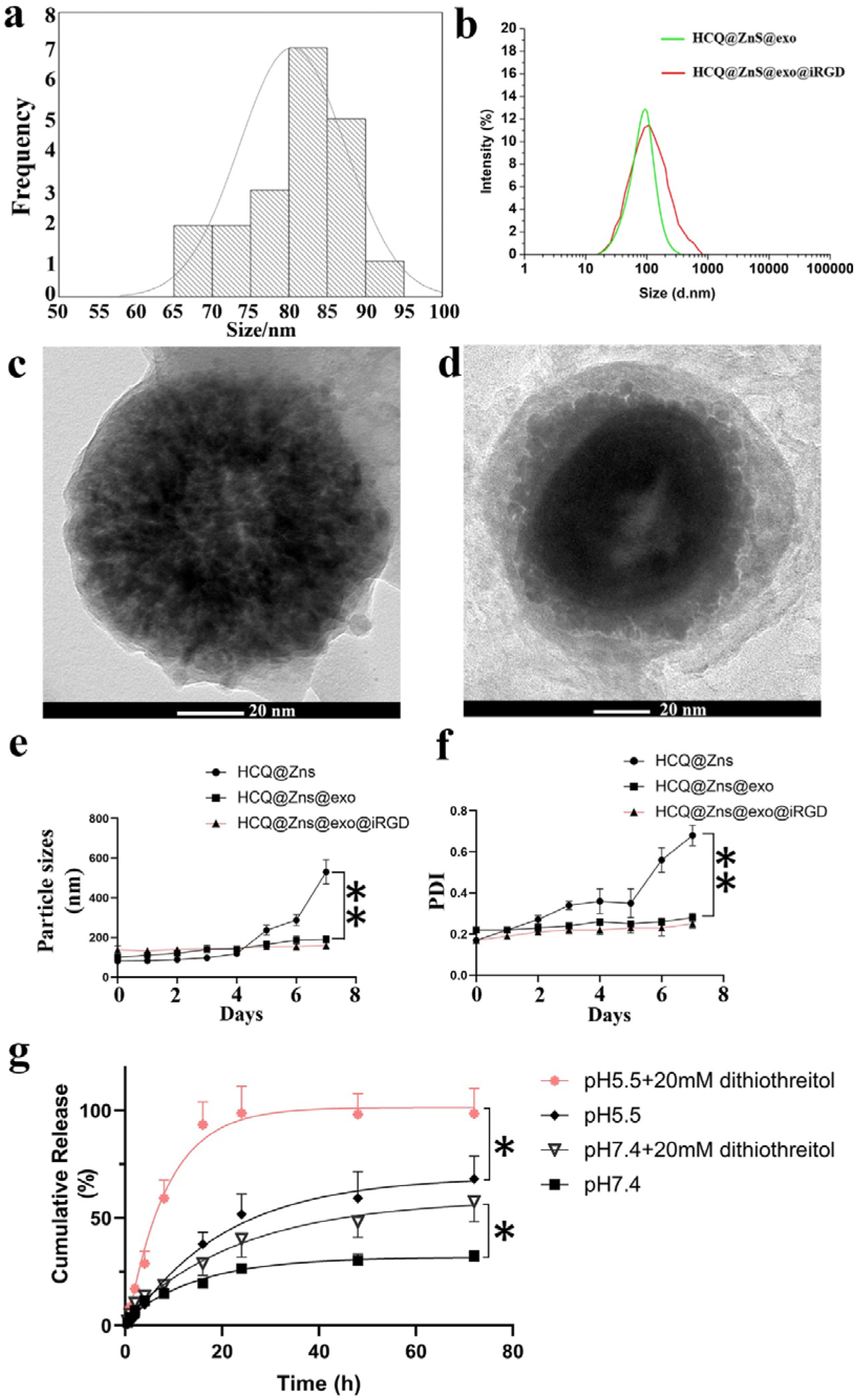
The ZnS based nanoparticles were characterized. The particle size distribution of **(a)** HCQ@ZnS, **(b)** HCQ@ZnS@exo and HCQ@ZnS@exo@iRGD. Transmission electron microscope images of **(c)** HCQ@ZnS@exo and **(d)** HCQ@ZnS@exo@iRGD. The stability of nanoparticles over 7 days in terms of **(e)** particle sizes and **(f)** PDI, respectively. **(g)** *In vitro* drug release study. The percentage release of hydroxychloroquine from different HCQ loaded ZnS based nanoformulations in various circumstances. Data shown are mean ± SD of 3 replicates. *, *p* < 0.05, **, *p* < 0.01.

TEM images showed a morphological characteristic of these nano-sized particles for HCQ@ZnS@exo (Figure 3c) and HCQ@ZnS@exo@iRGD (Figure 3d). These images of HCQ@ZnS@exo and HCQ@ZnS@exo@iRGD were compared for morphology, which exhibited a clear distinction between the nanoformulations. HCQ@ZnS@exo@iRGD with the dense network around the surface. Likewise, SEM images showed HCQ@ZnS@exo@iRGD (Figure S5b) have distinct different surface morphology with that of HCQ@ZnS@exo. There are brighter rims in HCQ@ZnS@exo@iRGD nanoparticles (Figure S5c).

To further examine the stability of these nanoparticles, we monitored their colloidal stability up to 7 ? days in terms of particle sizes and PDI, with the results depicted in Figure 3e and f. The particle size of HCQ@ZnS@exo and HCQ@ZnS@exo@iRGD remained constant during a 7-day incubation period. In contrast, the particle size of HCQ@ZnS increased dramatically after day six. (Figure 3e) Similarly, HCQ@ZnS@exo@iRGD and HCQ@ZnS@exo showed better stability relative to HCQ@ZnS. During the 7-day period, the PDI of HCQ@ZnS@exo@iRGD and HCQ@ZnS varied from 0.17 to 0.25 and 0.22 to 0.28 compared to 0.17 to 0.68 for HCQ@ZnS. (Figure 3f) These data show that, incorporation of the lipid layer of the liposome with the exosome in HCQ@ZnS resulted in enhanced stability of the engineered HCQ@ZnS@exo@iRGD.

### 3.4. Hydroxychloroquine loading and releasing study

The drug loading and releasing kinetics of ZnS@exo@iRGD were also investigated as depicted in *Supplementary information*. The encapsulation efficiency (EE) and drug loading (DL) of hydroxychloroquine in the different formulations are listed in Table 1. EE and DL of HCQ@ZnS were determined to be 27.10±2.21% and 15.20±2.31%, respectively. Almost no hydroxychloroquine leakage from ZnS was observed, as EEs of HCQ@ZnS@exo and HCQ@ZnS@exo@iRGD are as high as 97.5%.

With the assurance of good stability of HCQ@ZnS@exo@iRGD, we further studied drug release to explore the release kinetics at normal physiological pH (pH 7.4, PBS) and acidic (pH 5.5, acetate buffer) as depicted in Figure 3g. HCQ@ZnS@exo@iRGD showed similar bursts release in both pH conditions up to the first 8□h, with enhanced drug release characteristic in acidic pH compared to physiological pH when evaluated for further 36 h. Besides pH sensitive feature, higher release from HCQ@ZnS@exo@iRGD was also found under high GSH concentrations, which may attribute to redox-cleavable bond at outer layers of HCQ@ZnS@exo@iRGD. Overlapping these two factors together would lead to much steeper release curves. As shown in the red curve in Figure 3g, the accumulative release of hydroxychloroquine will reach plateau of 95% within just 24 h.

Overall, HCQ@ZnS@exo@iRGD showed a precisely controlled trend of drug release characteristic signifying that this engineering nanoparticles may be applied to complex tumor microenvironments.

### 3.5. Validation of fusion of lipid shell

Förster resonance energy transfer (FRET) and protein assay were carried out for the confirmation of fusion between the liposome and the hybrid exosomes. Lipids containing one of the FRET partners were incorporated within the liposomes. The change in the energy transfer with the fusion of the exosomes and liposomes can be seen in the change in the fluorescence spectra before and after fusion as shown in Figure 4a. The fluorescent intensity decreased after fusion. A diminished FRET signal after fusion with the exosomes suggests the distance between the FRET pair increases,[28] as expected when more lipids are incorporated in the bilayer as the liposomes and exosomes fuse.

**Figure 4.**
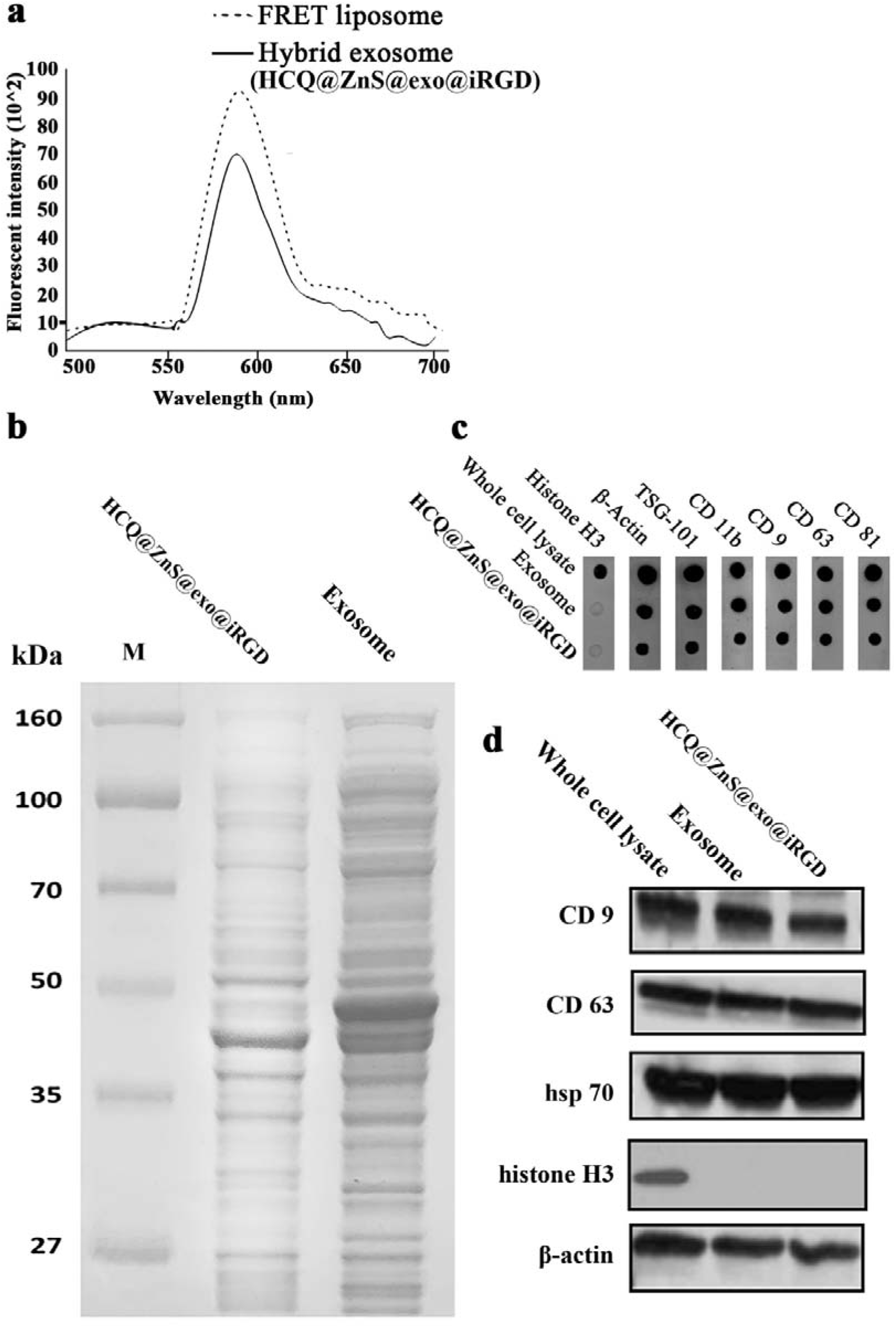
Validation of hybrid exosome of HCQ@ZnS@exo@iRGD formation. **(a)** Förster resonance energy transfer (FRET) study showing successful fusion of hybrid exosome (HCQ@ZnS@exo@iRGD) and liposome with FRET pairs. FRET study was conducted using fluorescent donor NBD (λem□=□525 □nm) and fluorescent acceptor RhB (λem□=D595□nm) at excitation wavelength of 470□nm; **(b)** SDS-PAGE analysis of exosome and HCQ@ZnS@exo@iRGD. Both samples were concentrated to achieve distinct protein bands; **(c)** the dot blot assay, and **(d)** the Western blot assay for the identification of exosome marker proteins in whole cell lysate, small extracellular vesicle (exosome), and hybrid exosome (HCQ@ZnS@exo@iRGD). β-actin was used as a positive control and histone H3 was used as negative control.

The protein cargo of the exosomes is important for their unique characteristics of homing to homogenous cells. As shown in Figure 4b, distinct protein bands in both exosome and hybrid exosome (HCQ@ZnS@exo@iRGD) were revealed in SDS-PAGE analysis after lipid fusion. Hybrid exosome showed similar protein bands to those observed with exosomes. More importantly, dot-blot assay showed the presence of major exosomal protein markers like transmembrane proteins (CD81, CD63, CD11b, and CD9) and tumor susceptibility gene 101 protein (TSG101) (Figure 4c).[29] These proteins were present in both the exosomes and the HCQ@ZnS@exo@iRGD, suggesting successful retention of major exosome proteins through the fusion process in HCQ@ZnS@exo@iRGD.

To further confirm the presence of exosome marker proteins in exosomes and HCQ@ZnS@exo@iRGD, the typical exosome marker proteins was analyzed with a Western blot assay. [30] As shown in Figure 4d, the Western blot assay confirmed the presence of exosomal markers CD9, CD63 and hsp70 in the HCQ@ZnS@exo@iRGD. [31] The protein β-actin was used as the positive control and histone H3, a nuclear protein, was used as a negative control. All three samples showed the presence of β-actin. However, the nuclear protein histone H3 was not present in exosome and HCQ@ZnS@exo@iRGD, which suggests that the isolated exosomes are free from possible nuclear contamination. These analyses further support the successful fabrication of HCQ@ZnS@exo@iRGD with the conservation of characteristic exosome protein cargoes.

### 3.6. Cellular internalization study

Cellular internalization of the Rh-B@ZnS@exo@iRGD system was studied with human U87 glioblastoma cells and human MCF10A breast epithelial cells using confocal imaging. Confocal images and quantitative analyses of the uptake behavior of the two cell lines showed the preferential internalization of different Rhodamine-B (Rh-B) labeled formulations after 3□h of treatment (Figure 5a and b). Significantly higher cellular internalization of Rh-B@ZnS@exo and Rh-B@ZnS@exo@iRGD was observed compared to Rh-B@ZnS in exosome parent cells U87, whereas there was no significant uptake difference among treatment groups in MCF10A cells. (Figure 5c and d) Different amount of Rh-B@ZnS@exo and Rh-B@ZnS@exo@iRGD internalized by U87 and MCF10A cells suggested uptake processes of Rh-B@ZnS@exo and Rh-B@ZnS@exo@iRGD in U87 are active transportation, while in MCF10A they are passive diffusion.

**Figure 5.**
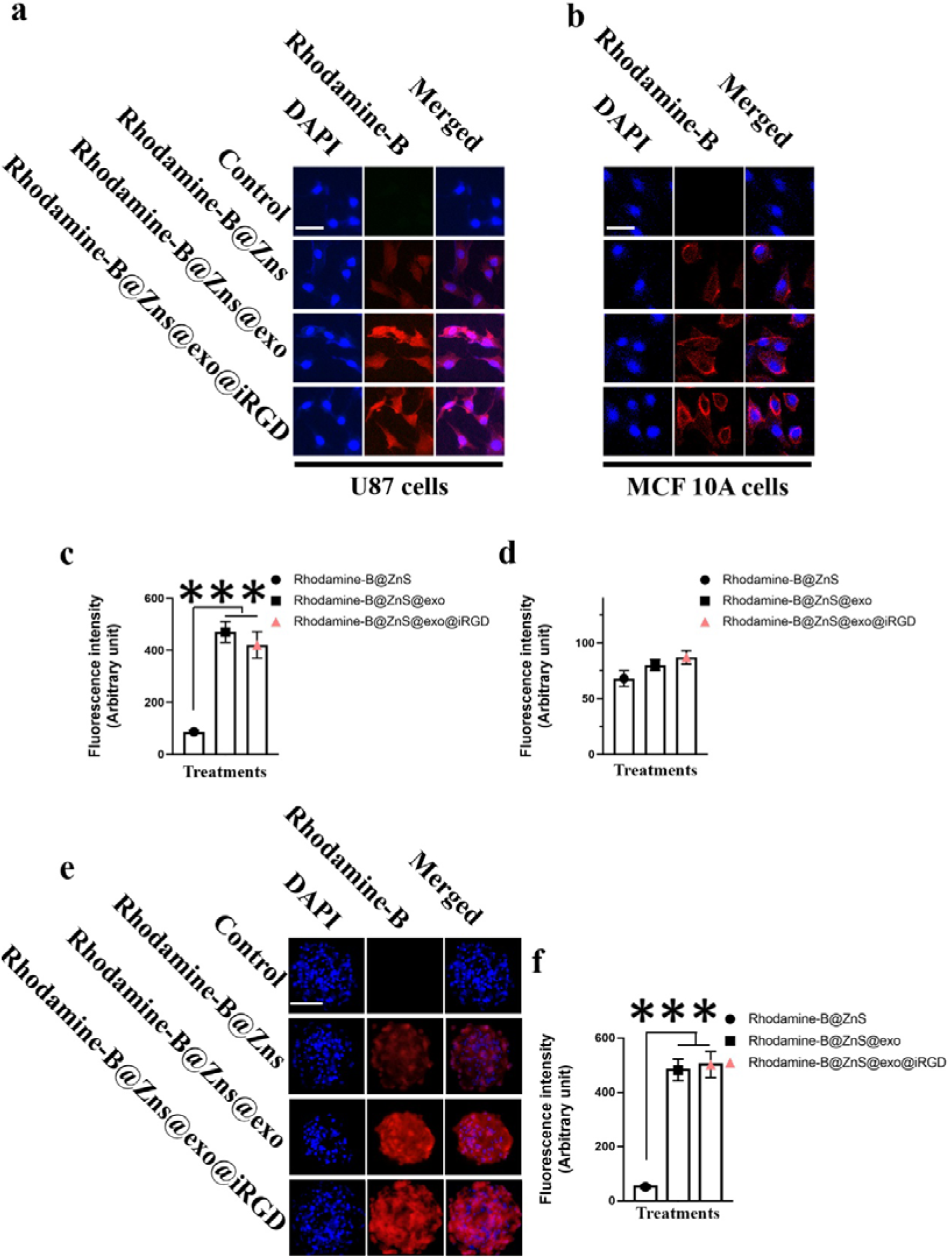
Cellular uptake behavior of the Rhodamine-B loaded different ZnS nanoparticles. Cellular uptake of different Rhodamine-B formulations in monolayer of **(a)** U87 and **(b)** MCF10A cells observed *via* confocal microscopy, scale bar 30 μm. Comparison of fluorescent intensities in **(c)** U87 and **(d)** MCF10A cells. *In vitro* **(e)** uptake behavior and **(f)** quantification of different Rhodamine-B loaded ZnS nanoparticles in U87 tumor spheroids, scale bar 100 μm. Internalization was quantified in terms of mean fluorescent intensity per tumor spheroid using Image-Pro Plus 7 software.[33] Data shown are mean ± SD of 3 replicates. ***, *p* < 0.001. Control group is without treatment.

To better understand the uptake behavior of the nanoparticles with a solid tumour,[32] 3D tumor spheroids were prepared. The uptake of the Rh-B@ZnS system in U87 tumor spheroids was low as determined by the weak fluorescence signal, indicating Rh-B@ZnS had difficulties in diffusing into the tumor spheroid. In contrast, the fluorescence intensity of Rh-B@ZnS@exo and Rh-B@ZnS@exo@iRGD are much higher, indicating that particles with exosome or hybrid exosome may help uptake by the tumor spheroids (Figure 5e). Quantitative measurements of the fluorescence intensity of U87 tumor spheroids confirmed the uptake efficiency of Rh-B@ZnS@exo and Rh-B@ZnS@exo@iRGD (Figure 5f) and this is consistent with the results shown in Figure 5e.

### 3.7. HCQ@ZnS@exo@iRGD decreases viability of U87 cells by lysosome de-acidification and autophagic flux blockage

To verify the cytotoxic effects of HCQ@ZnS@exo@iRGD, the *in vitro* anti-tumor effects of free drugs and drug-loaded nanoparticles against U87 cells were tested using MTT assay. A significant increase in the percentage of dead U87 cells was observed when treated with HCQ@ZnS@exo (IC_50_=36.5±4.8 μM), compared with those treated with free HCQ (IC_50_=75.9±12.2 μM) or HCQ@ZnS (IC_50_=68.6±3.5 μM). Likewise, the number of viable cells decreased significantly upon the treatment with HCQ@ZnS@exo@iRGD (IC_50_=34.9±3.1 μM). Even greater cytotoxic effects were observed when U87 cells were treated with HCQ@ZnS@exo@iRGD under light illumination (IC_50_=12.6±1.9 μM) (Figure 6a).

**Figure 6.**
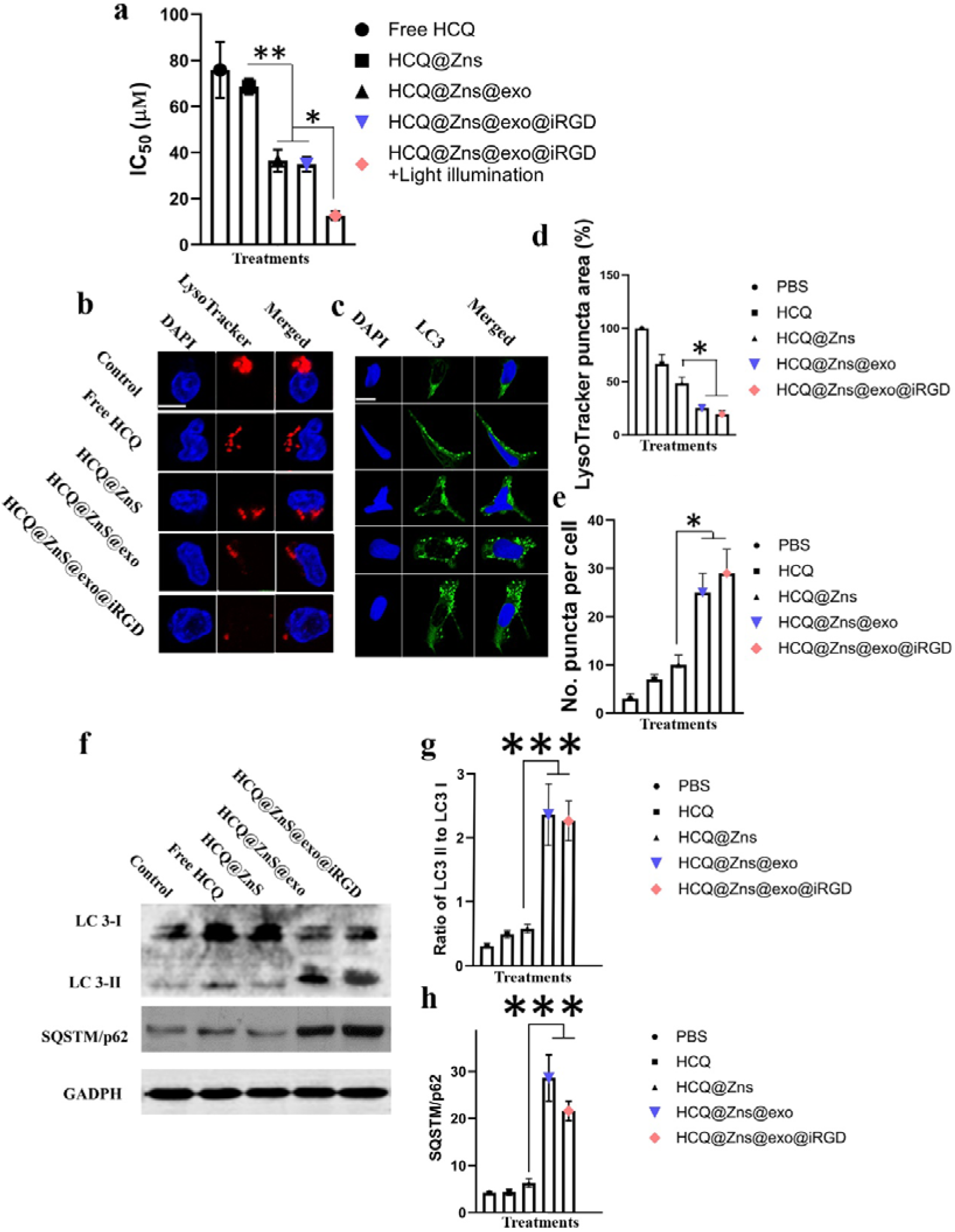
Cell viabilities of U87 cells after treatment with different formulations *in vitro*. **(a)** The IC50s for different formulations under different circumstance. U87 cells deactivate autophagy upon HCQ@ZnS@exo@iRGD administration. Representative images are shown for **(b)** Fluorescent staining of lysosomal (LysoTracker, red) and **(c)** autophagic markers (LC3, green) of U87 cells after treated with free HCQ, HCQ@ZnS, HCQ@ZnS@exo or HCQ@ZnS@exo@iRGD; Scale bar 10 ? μm in **b** and **c**. **(d)** The LysoTracker puncta area quantification using Image-Pro Plus 7 software and **(e)** the average number of LC3 puncta per cell (mean ± SD, *m2*= ? 30-45 cells) after different treatments. **(f)** Western blot for LC3 and SQSTM1/p62. **(g)** Quantification of the ratio of LC3-II to LC3-I expression using ImageJ software. **(h)** Quantification of levels of SQSTM1/p62 normalized to those in PBS-treated cells using ImageJ software. Data shown are mean ± SD of 3 replicates. *P < 0.05, **P < 0.01, ***, *p* < 0.001, according to a Student’s t-test.

Since hydroxychloroquine inhibits autophagy by blocking lysosomal acidification, we treated cells with free HCQ, HCQ@ZnS, HCQ@ZnS@exo or HCQ@ZnS@exo@iRGD at pH 7.4 and analyzed late endosomal compartments and lysosomes by staining with LysoTracker Red to visualize this process. U87 cells treated with HCQ@ZnS@exo and HCQ@ZnS@exo@iRGD showed less lysosomal red puncta area than other treatment groups (Figure 6b and d). To further compare the ability of free HCQ, HCQ@ZnS, HCQ@ZnS@exo or HCQ@ZnS@exo@iRGD to inhibit autophagy, an investigation of the occurrence of autophagy in U87 cells was performed by analyzing the expression pattern of microtubule-associated protein 1 A/1B-light chain 3 (MAP1LC3, also known as LC3).[34] Consistent with results of lysosome de-acidification, U87 cells showed increased expression of LC3 over time with more autophagosome puncta accumulated with HCQ@ZnS@exo or HCQ@ZnS@exo@iRGD treatment than with free HCQ or HCQ@ZnS treatment (Figure 6c and e).

As a further evaluation of the inhibition of autophagy, levels of LC3 and SQSTM1/p62, both of which play key roles in autophagy, were measured using Western blotting (Figure 6f). Both HCQ@ZnS@exo and HCQ@ZnS@exo@iRGD caused a larger increase in the ratio of LC3-II to LC3-I than free HCQ or HCQ@Zns after a 3 h treatment at a dose of 40 μM HCQ (Figure 6g), indicating greater inhibition. Consistently, SQSTM1/p62 expression was also higher with HCQ@ZnS@exo and HCQ@ZnS@exo@iRGD than the other treatments, confirming stronger autophagy inhibition (Figure 6h).

### 3.8. Biodistribution of nanoparticles *in vivo*

The orthotopic glioblastoma xenograft model was established for the assessments of the targeting ability of Cy 5.5@ZnS@exo@iRGD *in vivo* via IVIS Spectrum imaging system. Results show that the Cy 5.5@ZnS@exo@iRGD displayed a more selective accumulation at the glioblastoma sites 24 h post injection, compared with the Cy 5.5@ZnS@exo, Cy 5.5@ZnS or free Cy 5.5 (Figure 7a). After 24 h, the mice were sacrificed, major organs were excised for *ex vivo* imaging to display the tissue distribution (Figure 7b). Less nonspecific distribution of Cy 5.5@ZnS@exo@iRGD to major organs (except brain) was observed as the fluorescent intensities were much lower compared with other treatment groups. Moreover, the glioblastoma-bearing brains were also excised for *ex vivo* imaging, and the quantitative average radiant efficiency of glioblastoma-bearing brains also confirmed the highest glioblastoma aggregation of Cy 5.5@ZnS@exo@iRGD than the Cy 5.5@ZnS@exo, Cy 5.5@ZnS and free Cy 5.5 (Figure 7b). All results indicated Cy 5.5@ZnS@exo@iRGD held the highest glioblastoma targeting effect among different treatments, implying that the Cy 5.5@ZnS@exo@iRGD could easily cross the BBB and then uptake by the glioblastoma cells.

**Figure 7.**
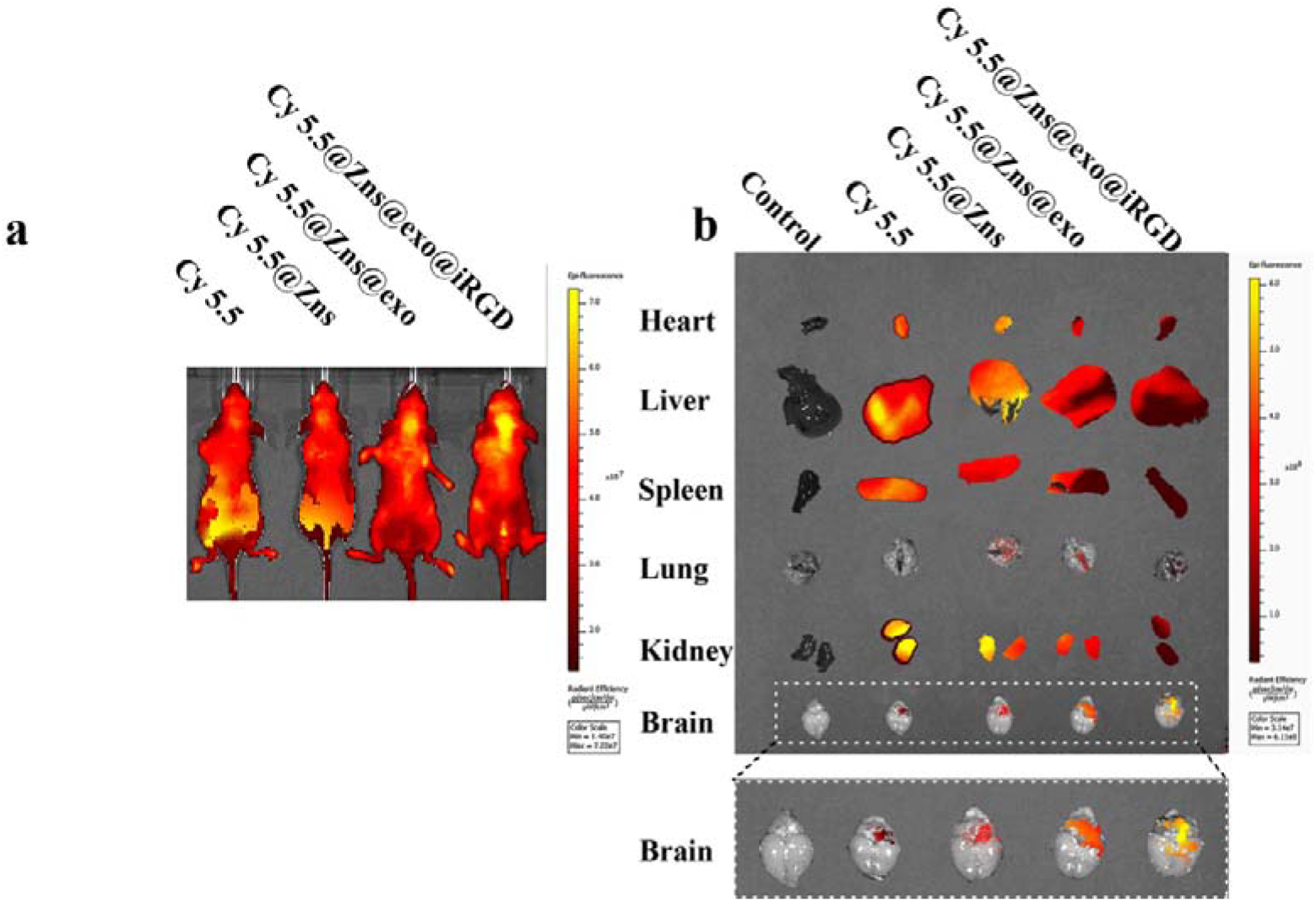
*In vivo* distribution of different Cy 5.5-loaded ZnS nanoparticles on the U87 orthotopic glioblastoma bearing mice model. **(a)** Whole body fluorescence imaging of U87 orthotopic glioblastoma mice at 24 h after tail vein injection of different Cy 5.5-loaded ZnS nanoparticles. **(b)** *Ex vivo* fluorescence imaging of the major organs and brain from orthotopic glioblastoma mice.

### 3.9. Anti-glioblastoma efficacy of nanoparticles in glioblastoma bearing mice

The anti-glioblastoma efficacy of nanoparticles was evaluated using the orthotopic glioblastoma (Luc-U87) mice model as illustrated in Figure 8a. The size of the glioblastoma in the brain were measured via the IVIS imaging system by intraperitoneal injection of D-luciferin potassium salt (150 mg/kg). As shown in Figure 8b, the HCQ@ZnS@exo@iRGD with light irradiation displayed obvious weaker luminescence intensity than other groups, which is consistent with the smallest area of luminescence due to their dual-targeted capability and powerful photodynamic therapy and efficient autophagic flux blocking as well. The glioblastoma growth curves determined by fluorescent intensity are illustrated in Figure 8c, showing much steeper curves for free HCQ and HCQ@ZnS treatment groups, while treatment with HCQ@ZnS@exo@iRGD under light illumination has the best efficacy in eliminating glioblastoma.

**Figure 8.**
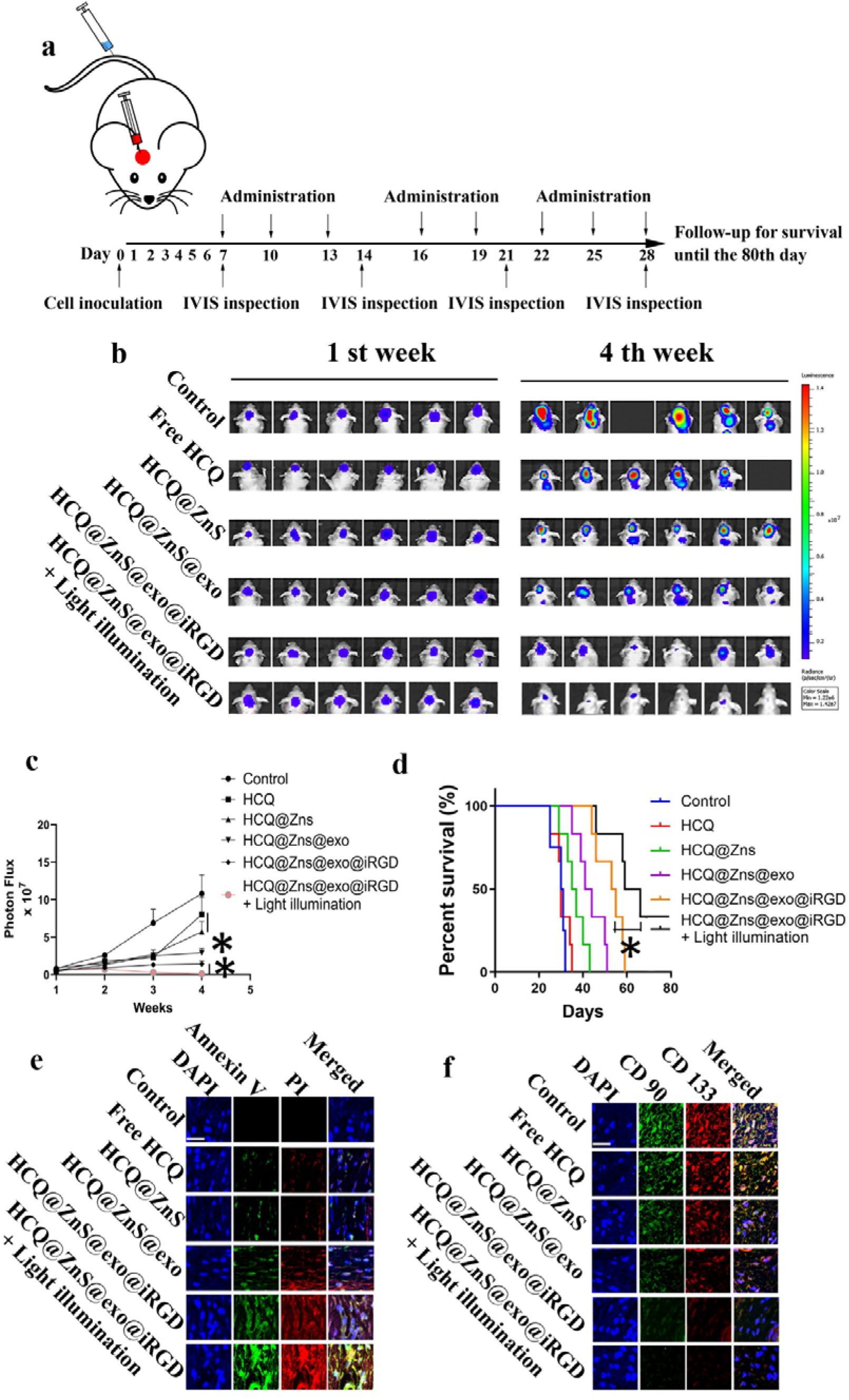
*In vivo* anti-glioblastoma efficacy of HCQ-loaded ZnS nanoparticles on the U87 orthotopic glioblastoma bearing mice model. **(a)** Schematic diagram of luciferase expressing orthotopic glioblastoma model and drug treatment. After glioblastoma cell implantation, tumor growth was monitored by IVIS imaging system. **(b)** Real-time bioluminescence images of the orthotopic glioblastoma model nude mice treated with PBS, free HCQ, HCQ@ZnS, HCQ@ZnS@exo or HCQ@ZnS@exo@iRGD with or without light irradiation on day of 7 and 28. The absence of mice in the imaging indicated the death of the mice. **(c)** Quantification of signals from the orthotopic glioblastoma bearing mice (*n* = 6, *\*p* < 0.05) on the day of 7, 14, 21 and 28. **(d)** Kaplan-Meier survival curves of the orthotopic glioblastoma nude mice (*n* = 6, \**p* < 0.05). **(e)** Staining of early apoptosis (annexin V) and late apoptosis (propidium iodide, PI) in tumors collected on day 14, scale bar 50 μm. (*n*=3) **(f)** Immunohistochemical staining of CD90 and CD133 in tumors from mice after the different treatments. HCQ@ZnS@exo@iRGD under light illumination significantly reduced expression of both markers, scale bar 50 μm. (*n*=3).

The therapeutic efficacy of clinical malignant tumor patients was mainly evaluated by overall survival time and life quality. The survival rate also confirmed improved efficacy of glioblastoma treatment by applying HCQ@ZnS@exo@iRGD with light irradiation. As shown in Figure 8d, of the mice treated with HCQ@ZnS@exo@iRGD and light irradiation displayed the longest median survival time (73 days), while PBS, free HCQ, HCQ@ZnS, HCQ@ZnS@exo or HCQ@ZnS@exo@iRGD without light irradiation had the median survival times of 30, 30, 36, 42.5 and 54 days, respectively.

Apoptosis at the glioblastoma sites after the different treatment conditions were measured by annexin V and PI staining. As shown in Figure 8e, the HCQ@ZnS@exo@iRGD with light irradiation group induced more cell apoptosis than the other groups. CD90 is a multi-structural and multi-functional cell surface receptor involved in cell proliferation, cell differentiation, cell migration and angiogenesis, while CD133 is believed to be associated with tumorigenicity and progression of glioblastoma.[35, 36] Both CD90 and CD133 are often used as surface markers of glioblastoma.[37, 38] The expression of CD90 and CD133 in cancer cells in tumors after treatment with the different formulations was investigated, *in vivo*. As shown in Figure 8f, significant reduced expression of CD90 and CD133 following treatment with HCQ@ZnS@exo@iRGD under light illumination was observed, suggesting that HCQ@ZnS@exo@iRGD was effective for targeting and destroying CD90^+^CD133^+^ glioblastoma in multiple ways. All these results suggest that HCQ@ZnS@exo@iRGD may efficiently deliver HCQ@ZnS to glioblastoma site for targeting and eliminating glioblastoma cells with light irradiation *in vivo*.

No obvious signs of side effects were observed after treatment with HCQ@ZnS@exo@iRGD. No deaths or significant loss of body weight occurred following treatment with HCQ@ZnS@exo@iRGD (Figure S7a). Hematological indices after treatment with HCQ@ZnS@exo@iRGD were like those of healthy animals (Figure S7b). These results suggest that the HCQ@ZnS@exo@iRGD shows minimal systemic toxicity *in vivo*.

## 4. Discussion

Glioblastoma is a devastating and deadly malignant primary brain tumor in adults. It nearly always relapses after initial conventional treatment, and frequently exhibits resistance to current therapeutics. [39] A possible reason for the recurrence and treatment resistance is the presence of the blood-brain barrier (BBB) and unfavourable tumor microenvironment (TME).[40] Therefore, targeted elimination of glioblastoma cells might be effective for glioblastoma eradication. [41] However, recent treatments for targeting glioblastoma cells have been less successful, as multiple major hurdles to glioblastoma cells exist.

The transportation of anti-tumor agents to cross the BBB remains one of the main obstacles for effective glioblastoma treatment.[42] Since the protein expressions of integrin αvβ3 and neuropilin-1 are relatively high in the endothelial and glioblastoma cells, grafting iRGD ligand on the surface of HCQ@ZnS@exo greatly increase the *in vivo* cellular uptake of HCQ@ZnS@exo@iRGD in the orthotopic glioblastoma mice model than those of ZnS nanoparticles without iRGD ligand modification. Moreover, pH- and redox-cleavable linker was designed and synthesized to connect iRGD with phosphatidylserine. The fastest release of hydroxychloroquine from HCQ@ZnS@exo@iRGD at pH 5.5 with high GSH suggests that the dual-stimuli responsive phosphatidylserine-*cis*-aconitic anhydride-cystamine-PEG-iRGD influence the releasing characteristic of the HCQ@ZnS@exo@iRGD. Under acidic conditions with high level of GSH, the aconitic anhydride bond and cystamine linker would rupture and change the structure of the HCQ@ZnS@exo@iRGD leading to differential release of hydroxychloroquine. The results were also consistent with the biodistribution of HCQ@ZnS@exo@iRGD *in vivo*, since the pH- and GSH-sensitive iRGD becomes cleaved in tumor microenvironment, where H^+^ and GSH concentration are rather high.

Exosomes are 30-100 nm extracellular vehicles (EVs) released by all living cells, including glioblastoma cells and glioblastoma cells.[43] These small vesicles contain lots of bioactive materials in the form of proteins, DNA, mRNA, miRNA and lipids, acting as information carriers, playing important roles in intercellular communication by transferring their molecular contents to recipient cells. Exosomes derived from glioblastoma cells carry membrane receptors and signaling proteins that participate in glioblastoma progression and purposely aim for glioblastoma cells.[44] In the present study, exosomes were isolated from human U87 glioblastoma spheroids cell culture media and characterized. The subsequently exposed exosome membrane layer showed clear evidence in specifically guiding the hydroxychloroquine loaded nanoparticles to glioblastoma cells due to homing ability of exosome. [45] Owing to the tailored double-targetabilities of HCQ@ZnS@exo@iRGD, the preferential accumulation of the particles in glioblastoma cells was observed *in vitro* and *in vivo*.

The surface morphology, particle size and ζ-potential are all important characteristics of drug loaded nanocarriers.[46] They have been demonstrated to play important roles in determining cellular and tissue uptake efficiency and toxic effect on cells. [47] Smooth surface morphology has been demonstrated to provide the uniformity and stability of the system. [48] The nanoparticle delivery vehicle we have produced has an obvious core-shell structure (Figure 3b), suggesting that the exosome membranes and iRGD-derived liposome decorate the surface of the HCQ@ZnS. The size and zeta potential of NPs not only impact on their colloidal stability but also influence the effectiveness of their interaction with negatively charged cell membranes, which is the pivotal step for successful BBB penetration and glioblastoma cells uptake. Unmodified HCQ@ZnS was shown to have a slight positive surface charge, due to existing cationic Zn^2+^. In contrast, the ζ-potential of HCQ@ZnS@exo and HCQ@ZnS@exo@iRGD decreased significantly, indicating that anionic exosome membrane and/or iRGD-derived liposome located on the surface of HCQ@ZnS neutralized the charge of the nanoparticles. The positive surface charge of ZnS core is necessary to ensure the uptake by targeted cells due to electrostatic interactions between negatively charged cellular membranes and positively charged nanocarriers.[49]

Photodynamic therapy (PDT) has shown promising effect in cancer treatment. [50] Metal materials including ZnS, Au and Pt are excellent therapeutic materials for phototherapies. Some of them have achieved a higher selectivity than the conventional treatments towards tumor sites as only the photosensitizer accumulated in the tumor sites can be activated by the light irradiation.[51] However, spontaneous elevated autophagic activities in tumor cells severely compromised PDT’s efficacy by digesting those impaired organelles to provide necessities for maintaining cell grows. [52] In this regard, the hydroxychloroquine loaded ZnS nanoparticles were synthesized to achieve the combination of photodynamic therapy and chemotherapy. All results confirmed a higher anti-tumor efficacy of HCQ@ZnS@exo@iRGD with visible light irradiation than the other groups both *in vitro* and *in vivo*.

The current studies aimed to determine whether autophagy inhibition has a beneficial role in glioblastoma treatments. As hydroxychloroquine is an FDA-approved drug for inhibiting autophagy, this compound has been extensively used to test whether blocking this pathway will improve tumor treatments. Our study underlines that hydroxychloroquine loaded into hybrid exosome covered ZnS nanoparticles can substantially impair the autophagic flux. As a result, positive effects on tumor regression in the orthotopic glioblastoma mice model by treatment with HCQ@ZnS@exo@iRGD was observed in our research.

It is widely accepted that hydroxychloroquine accumulate in lysosomes (lysosomotropism) and inhibit their function leading to accumulation of autophagic vesicles in tumor cells.[53] However, its clinical relevance has been severely limited as high doses are required to compensate for its nonselective distribution *in vivo* and its inefficiency to cross the BBB and further target glioblastoma cells.[54] Here, we avert all these issues by encapsulating hydroxychloroquine in hollow ZnS nanoparticles coated with hybrid exosome. Using *in vitro* and *in vivo* approaches we showed that HCQ@ZnS@exo@iRGD specifically targets glioblastoma and further efficiently accumulates hydroxychloroquine within glioblastoma cells. Autophagy inhibiting would intensely enhance the antitumor efficiency of photodynamic therapy. Therefore, the encapsulation of hydroxychloroquine into the ZnS@exo@iRGD nanoparticles can effectively combine photodynamic therapy with chemotherapy to show high efficiency for the treatment of glioblastoma.

The cellular uptake of HCQ@ZnS@exo@iRGD by MCF10A cells verified that the HCQ@ZnS@exo@iRGD effectively reduced the toxicity of hydroxychloroquine to normal cell by possible reason of exosome coating. Furthermore, data from *in vivo* distribution investigation also confirmed that encapsulating hydroxychloroquine in HCQ@ZnS@exo@iRGD significantly reduced its distribution to the kidney, where hydroxychloroquine elicits chronic renal failure.[55] All above results suggest selective, efficient internalization of HCQ@ZnS in glioblastoma cells and the resulting accumulation of hydroxychloroquine in lysosomes implies not only greater therapeutic effect but also lower systemic toxicity. Therefore, the HCQ@ZnS@exo@iRGD are promising alternative for therapeutic delivery to the glioblastoma cells.

## 5. Conclusion

In summary, we designed a multifunctional nanoplatform based on multi-shelled hydroxychloroquine loaded hollow ZnS spheres for photodynamic therapy/chemotherapy of glioblastoma. The so-called HCQ@ZnS@exo@iRGD possess the following features: (1) specific glioblastoma cells targeting, (2) efficient ROS generation under light irradiation, (3) superb blockage of autophagic flux by hydroxychloroquine, which enable combination of photodynamic therapy and chemotherapy. A good understanding of the *in vivo* behavior of HCQ@ZnS@exo@iRGD, such as their quantitative pharmacokinetics, subcellular distribution, and long-term toxicity, is of importance and requires further exploration. HCQ@ZnS@exo@iRGD opens a new therapeutic window for the treatment of glioblastoma. Therefore, HCQ@ZnS@exo@iRGD stands out as a highly potent and versatile therapeutic composite in the field of nanomedicine.

## Author contributions

J. Mo designed the research. J. Mo, M. Li, and S. Li carried out the experimental studies. J. Mo and W. Zhao interpreted the data and wrote the manuscript, which E. Tang and J. Gooding helped edit. All authors read and approved the manuscript.

## Acknowledgements

This project was supported by the Natural Science Foundation of Guangxi Province, (2019GXNSFAA245095). The authors thank the Mark Wainwright Analytical Centre at the University of New South Wales.

