## supplementary information for "Zinc Sulfide-Based Hybrid Exosome-Coated Nanoplatform for Targeted Treatment of Glioblastoma in an Orthotopic Mouse Glioblastoma Model" (from another version of this preprint: version 4, 2020-09-05)

**Organic Synthesis**
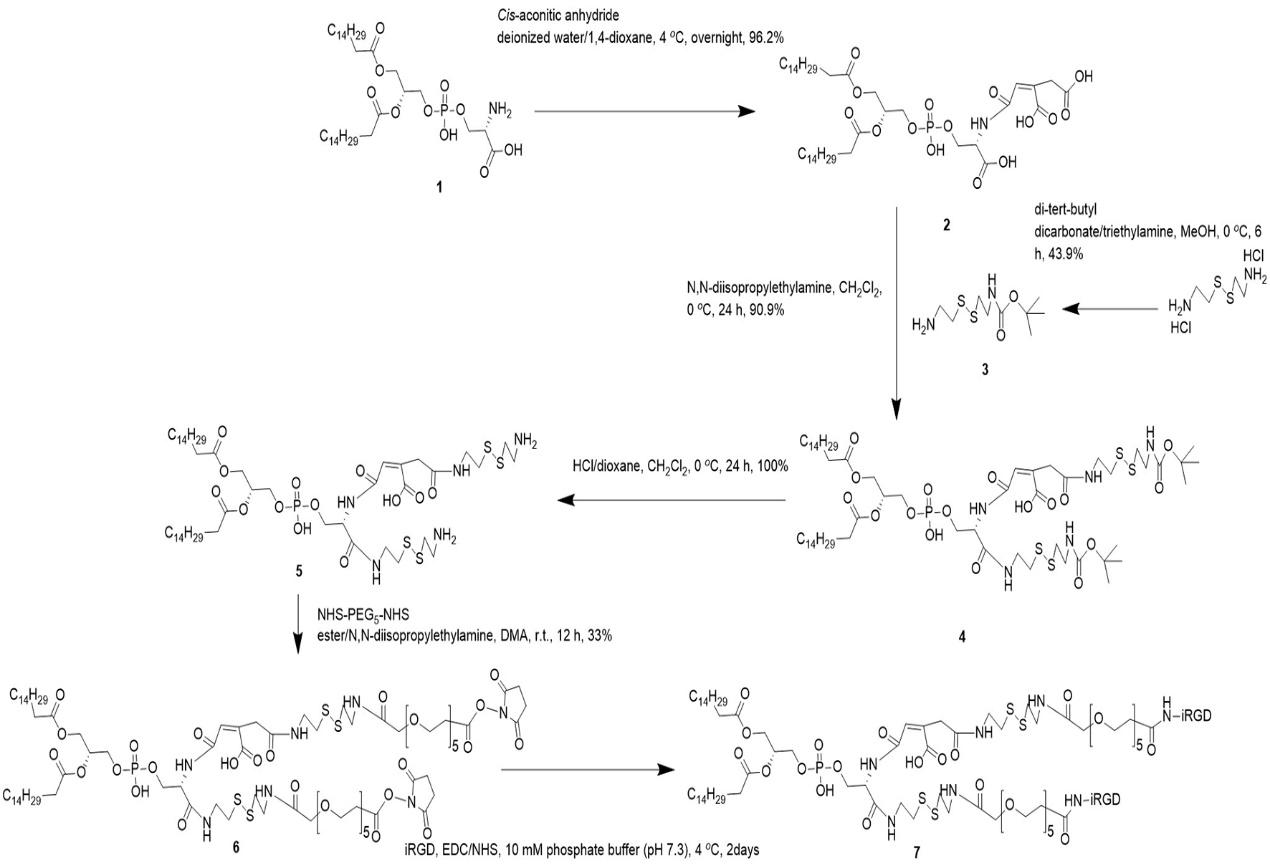

**Figure S1.** Graphical synthetic routes of phosphatidylserine-*cis*-aconitic anhydride-cystamine-PEG-iRGD.

Phosphatidylserine amidol succinic acid (**2**)

Phosphatidylserine (**1**, 10 g, 13.60 mmol) was dissolved in 4 L deionized water. *Cis*-aconitic anhydride (3.12 g, 20 mmol) in 100 mL of 1,4-dioxane was added slowly, and the mixture was stirred overnight at 4°C. The solution was mixed with 500 mL of chloroform and 5 mL of 5% w/v aqueous sodium bicarbonate. The chloroform phase was decanted and the residual solution extracted with ethyl acetate. The extract was concentrated on a rotary evaporator and dried at room temperature under vacuum to get **2** as yellow solid. Yield: 11.66 g (96.2%). ^1^H NMR (400 MHz, CDCl_3_): δ 5.34 (s, 1H, C=CHC=O), 5.58 (quint, 1H, CO-CH-CO), 1.12-1.34 (m, 6H, CH_3_).

N-tert-Butoxycarbonyl cystamine (**3**)

Cystamine dihydrochloride (6.756 g, 30.0 mmol) and triethylamine (12.54 mL, 90.0 mmol) were dissolved in MeOH (90 mL) and added to a methanolic solution (60 mL) of di-tert-butyl dicarbonate (3.273 g, 15.00 mmol). The reaction mixture was stirred for 6 h at 0°C, after which the solution was evaporated, and 1 M NaH_2_PO_4_ was added to the residue. The aqueous solution was washed with diethyl ether to remove di-t-Boc-cystamine. The aqueous solution was adjusted to pH 9 by addition of 1 M NaOH, and the solution was extracted with AcOEt. The organic phase was dried over MgSO_4_ and then evaporated. The resulting yellow oil was dried in vacuo to give **3**. Yield: 1.661 g (43.9%). ^1^H NMR (400 MHz, CDCl_3_): δ 3.66 (t, 2H, C(=O)-NHCH_2_), 3.33 (t, 2H, CH_2_NH), 2.89 (t, 4H, CH_2_SSCH_2_), 1.56 (s, 9H, CH_3_).

Phosphatidylserine-*cis*-aconityl-PEG-ss-Boc (**4**)

Compound **2** (2.674 g, 3 mmol) was conjugated to compound 3 in an EDC/NHS coupling reaction. Compound 3 (2 g, 7.92 mmol) was dissolved in distilled water (200 mL), followed by dilution with 200 mL methanol. A predetermined amount of compound **2** was dissolved in 200 mL of dimethylformamide, then equal amounts of EDC and NHS (two equivalents of carboxyl group) were added to the solution. After 15 min of stirring, the solution obtained above was added. The resulting solution was stirred for 1 day at room temperature. The solution was dialyzed against excess distilled water. The product was freeze-dried to obtain **4** as yellow solid. Yield: 3.711 g (90.9%). ^1^H NMR (400 MHz, CDCl_3_): δ 6.25 (s, 1H, C=CHC=O), 5.53 (quint., 1H, CO-CH-CO), 4.56 (d, 6H, O-CH_2_CH) 3.62 (t, 2H, C(=O)-NHCH_2_), 3.40 (t, 8H, CH_2_NH), 2.93 (s, 1H, C(=O)-CHC), 2.85 (t, 8H, CH_2_SSCH_2_), 1.52 (s, 9H, CH_3_), 1.15-1.31 (m, 6H, CH_3_).

N-Methacryloylcystamine (**5**)

Compound **4** (1.36 g, 1 mmol), dissolved in CH_2_Cl_2_ (100 mL), was added to 4 N HCl/dioxane (1.5 mL, 5.96 mmol) in CH_2_Cl_2_ (100 mL), and stirred for 24 h at 0°C. The solution was evaporated to give the crude product **5** as a white solid, which was washed with diethyl ether and dried in vacuo. Yield: 1.116 g (96.2%). ^1^H NMR (400 MHz, CDCl_3_): δ 6.33 (s, 1H, C=CHC=O), 5.53 (quint., 1H, CH_2_O-CH-CH_2_O), 4.56 (d, 6H, O-CH_2_CH) 3.62 (t, 2H, C(=O)-NHCH_2_), 3.40 (t, 8H, CH_2_NH), 2.93 (s, 1H, C(=O)-CHC), 2.85 (t, 8H, CH_2_SSCH_2_), 1.15-1.31 (m, 6H, CH_3_).

NHS-type cleavable monomer for conjugated template (**6**)

Compound **5** (0.621 g, 0.535 mmol) and N,N-diisopropylethylamine (0.095 mL) were dissolved in DMA (100 mL), after which NHS-PEG_5_-NHS (0.605 g, 1.2 mmol) was added to the solution. The reaction mixture was stirred for 12 h at room temperature and evaporated in vacuo. The residue was dissolved in CHCl_3_ and washed with brine, and the organic phase was dried over MgSO_4_ and evaporated. The resulting yellow solid was dried in vacuo to give **6**. Yield: 0.342 g (33.0%). ^1^H NMR (400 MHz, CDCl_3_): δδ 6.28 (s, 1H, C=CHC=O), 5.53 (quint., 1H, CH_2_O-CH-CH_2_O), 4.53 (d, 6H, O-CH_2_CH) 3.62 (t, 2H, C(=O)-NHCH_2_), 3.52 (t, 20H, NHC(=O)-CH_2_(CH_2_OCH_2_)_5_), 3.41 (t, 8H, CH_2_NH), 2.95 (s, 1H, C(=O)-CHC), 2.83 (t, 8H, CH_2_SSCH_2_), 2.55 (t, 8H, C(=O)CH_2_CH_2_C(=O)), 1.17-1.30 (m, 6H, CH_3_).

Synthesis of template molecule phosphatidylserine-*cis*-aconitic anhydride-cystamine-PEG-iRGD (**7**)

Compound **6** (0.194 g, 0.10 mmol), EDC (0.50 mmol) and NHS (0.50 mmol) were dissolved into phosphate buffered solution (PBS) (pH = 6.0) and stirred at room temperature for 4 h to activate the carboxyl group of compound **6**. Then, iRGD peptide (0.284 g, 0.3 mmol) was added to the above solution. The above mixture was stirred at room temperature for 48 h. The solution was subsequently dialyzed with a dialysis tubing (Fisher Scientific, MWCO: 3.5 kD) to remove unreacted monomers. The dialyzed solution was ultra-filtered using an Amicon® Ultra-15 (4000 rpm, 10 min) and lyophilized to give **7** as a white solid, Yield: 0.362 g (100 %).

**Characterizations of phosphatidylserine-*cis*-aconitic anhydride-cystamine-PEG-iRGD**

The phosphatidylserine-*cis*-aconitic anhydride-cystamine-PEG-iRGD compound was first characterized using ^1^H NMR spectrum in CDCl_3_. The spectra were in accordance with the expected structures. As shown in Figure S2a, the signals at 5.08 ppm (peak b) corresponded to the ethenyl protons in phosphatidylserine, which indicated that PS (phosphatidylserine) was successfully activated. The signals at 6.37 ppm (peak a) corresponded to the methylene protons in aconitic anhydride as acid cleavable linker. Protons in PEG chain was observed at 3.45 ppm (peak d). Finally, peak c (δ=5.30 ppm) was allocated as protons in characteristic methine of iRGD.

The respective characteristic peaks of PEG chain, phosphatidylserine and iRGD peptides were also observed in the FTIR spectra of final product of phosphatidylserine-*cis*-aconitic anhydride-cystamine-PEG-iRGD (Figure S2b). Specifically, PEG shows characteristic stretching frequencies for -CH_3_ symmetric, and C-O, at 2868 and 1061 cm^−1^, respectively. The peaks at 1672 and 1540 cm^-1^ were assigned to amide bands from iRGD. After conjugating iRGD with PEG cystamine *cis*-aconitic anhydride phosphatidylserine (PEG-ss-aPS), the obvious characteristic amide bands with a slight blue shift (1625 and 1503 cm^-1^) with more intense absorption bands also being observed. Together these indicate the successful conjugation of acid- and redox-cleavable PEGylated phosphatidylserine with iRGD by the amide bond. Furthermore, the peak at 1063 cm^-1^ characteristic of the υ_3_ band of HPO_4_^2-^ are observed in both spectrum of phosphatidylserine and phosphatidylserine-*cis*-aconitic anhydride-cystamine-PEG-iRGD. Therefore, the designated product as phosphatidylserine-*cis*-aconitic anhydride-cystamine-PEG-iRGD was successfully synthesized, as demonstrated in the FTIR spectra.

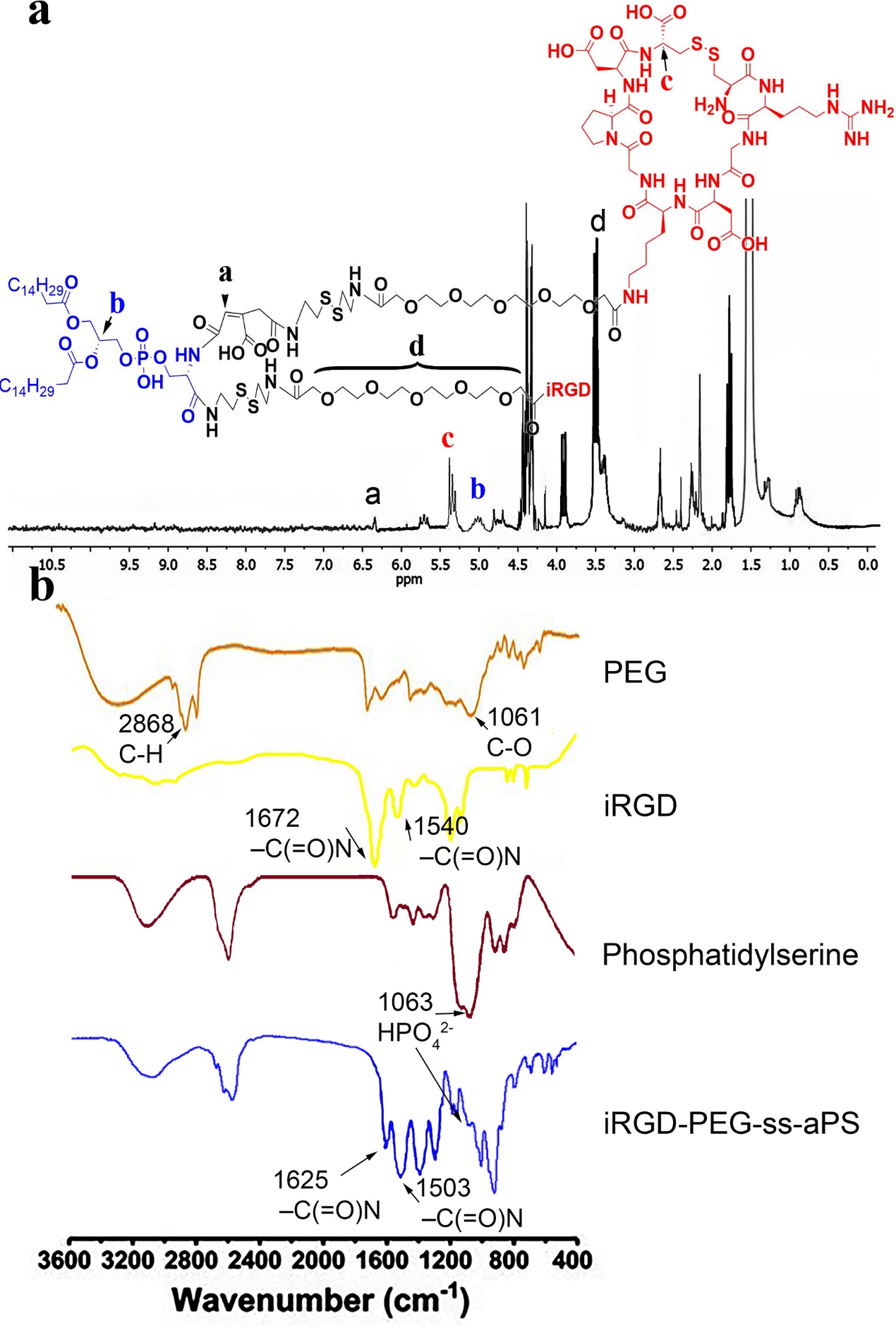

**Figure S2.** Chemical structure of phosphatidylserine-*cis*-aconitic anhydride-cystamine-PEG-iRGD (iRGD-PEG-ss-aPS) was confirmed by **(a)** ^1^H NMR and **(b)** FTIR spectrum.

**XPS survey spectrum of the ZnS nanoparticles**

The chemical state of the elements in ZnS sample was investigated by X-ray photoelectron spectroscopy (XPS). The wide XPS scan spectrum (Figure S3a) shows that except for Zn and S elements, the peaks of O, N, C elements can also be observed. The abundant N and C elements in ZnS should derive from surface-absorption species released by gelatin. As shown in Figure S3b, the binding energies (BE) of Zn2p_3/2_ and Zn2p_1/2_ peaks in ZnS are about 1021.4 eV and 1044.5 eV, respectively, which are in accordance with the BE values of Zn^2+^ in ZnS.^[1]^ Furthermore, the peaks of S2p_3/2_ and S2p_1/2_ located at 161.9 eV and 163.5 eV (Figure S3c), are consistent with S^2−^ in ZnS. The peak of N 1s can be deconvoluted into two peaks located at about 400.0 eV and 401.5 eV, which can be ascribed to the pyridine-*N* and quaternary-*N* determined by the structure of gelatin itself. (Figure S3d)

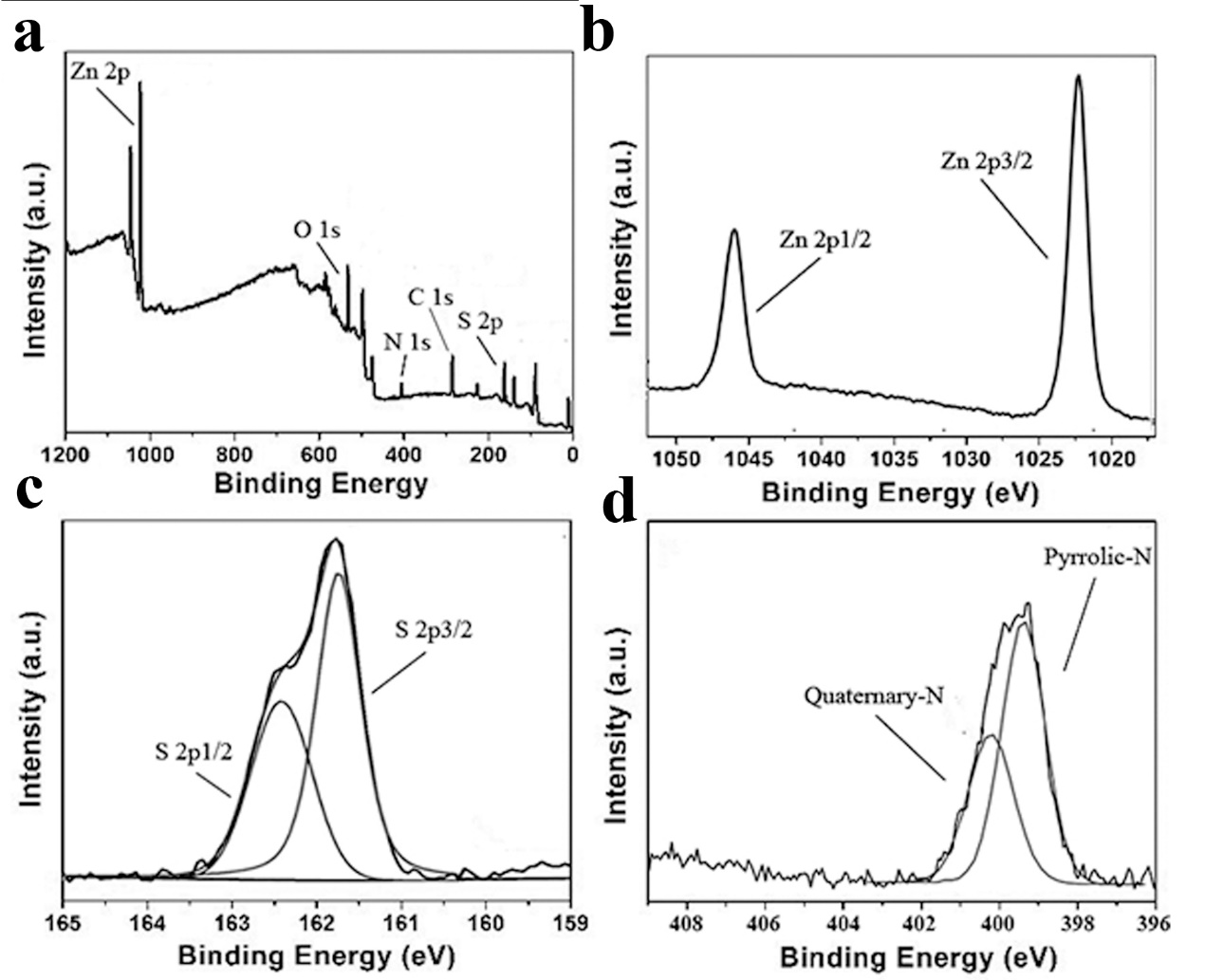

**Figure S3. (a)** The wide scan XPS spectrum and **(b)** Zn 2p, **(c)** S 2p and **(d)** N 1s spectra of ZnS sample.

**Photocatalytic activity**

The photocatalytic performances of ZnS nanoparticles were characterized by degradation of MB in different aqueous media under the irradiation of the 500 W tungsten halogen lamp for a specific period. Prior to illumination, the suspension was magnetically stirred for 30 min in the dark to achieve the adsorption equilibrium of the MB on the photocatalyst powders. The absorption spectra of MB solutions in the presence of ZnS sample under different test conditions is shown in Figure S4.

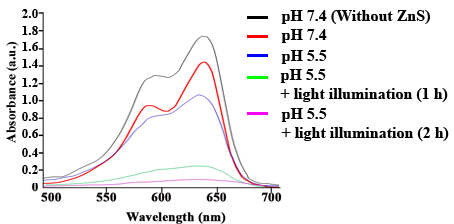

**Figure S4.** Absorption spectra as of MB solution obtained with/out the presence of ZnS in different test conditions.

**Exosome purification and hybridization with synthetic liposome**

Exosomes were isolated from GSC conditional media by using total exosome isolation (from cell culture media) reagent (Invitrogen), according to the manufacturer’s instruction.^[2]^ Exosomal protein was quantified using the Pierce™ BCA protein assay kit (ThermoFisher). The morphology of exosomes was observed by transmission electron microscopy, and the marker expressions were evaluated by Western blot.

Harvested sEVs were hybridized with synthetic liposomes using thin film hydration technique followed by extrusion through 200 nm size polycarbonate membranes. Synthetic liposome was prepared using phosphatidylserine:phosphatidylserine-*cis*-aconitic anhydride-cystamine-PEG-iRGD (molar ratio of 5:1) and cholesterol in a molar ratio of 70:30 following thin film hydration technique. The ratio of the liposome to sEV was optimized to 5:1. sEVs was quantified based upon protein content whereas liposome was quantified based upon lipid weights. 200 µg protein equivalent of sEVs dispersed in 1 mL Phosphate buffer saline (PBS) was added to 600 µg of lipid film to form hybrid exosome after extrusion by 200 nm polycarbonate membrane filter.

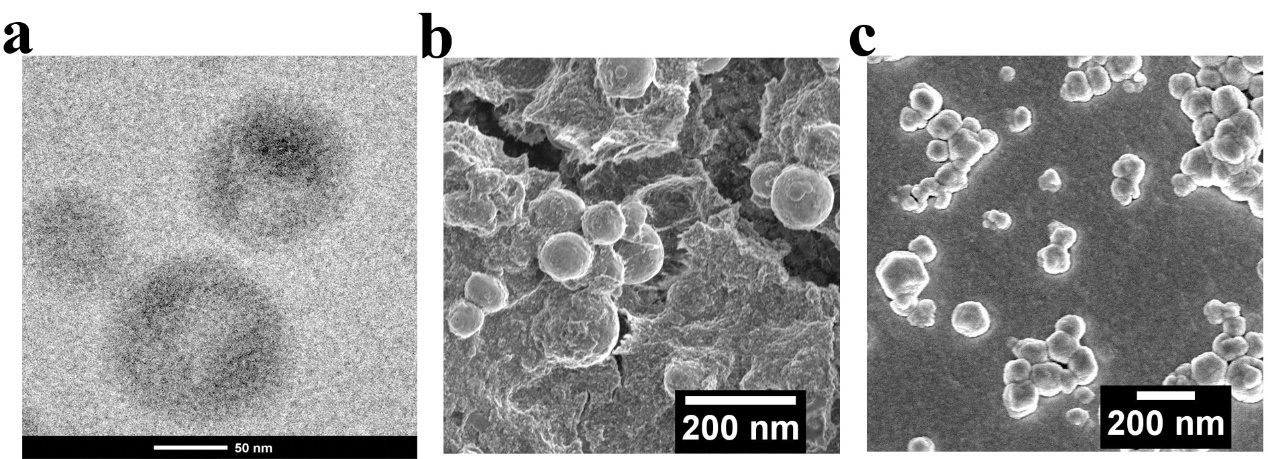

**Figure S5.** Transmission electron microscope images of **(a)** purified exosome (exo) and scanning electron microscope images of **(b)** HCQ@ZnS@exo and **(c)** HCQ@ZnS@exo@iRGD.

**LC-MS/MS conditions**

Liquid chromatography was performed on a Shimadzu HPLC system (Shimadzu, Tokyo, JPN). Hydroxychloroquine and I.S. (hesperetin) were separated on a 50 × 2.1 mm BetaBasic C8, 5 μm column (Thermo Hypersil-Keystone, Bellefonte, USA). The mobile phase comprising 0.1% v/v formic acid-acetonitrile (94:6, v/v) was delivered isocratically at a flow rate of 0.5 mL/min. Hydroxychloroquine and the I.S. were monitored using a Finnigan TSQ Quantum EMR Triple Quadrupole mass spectrometer (Thermo Fisher Scientific Corp., San Jose, USA) equipped with an electrospray ionization (ESI) source. The mass spectrometer was operated in positive ESI mode with a collision gas (Argon) pressure of 1.5 mTorr, a typical electrospray needle voltage of 3000 V, a sheath nitrogen gas flow of 49 (arbitrary unit) and a heated capillary temperature of 350°C. Hydroxychloroquine was analyzed by the multiple reaction monitor (MRM) mode using ion transitions at a proper collision energy as follows: The precursor to product ion transitions of m/z 336 → 247 and m/z 340 → 251 were used to measure the analyte and IS, respectively. The mass spectrometer was tuned to its optimal sensitivity by direct infusion of Hydroxychloroquine. The assay demonstrated a good linear dynamic range of 1-3200 ng/mL for hydroxychloroquine, with coefficient of determination (r^2^) of 0.997. The chromatographic run time was 8 min. The retention time of hydroxychloroquine is 5.28 min. (Figure S6)

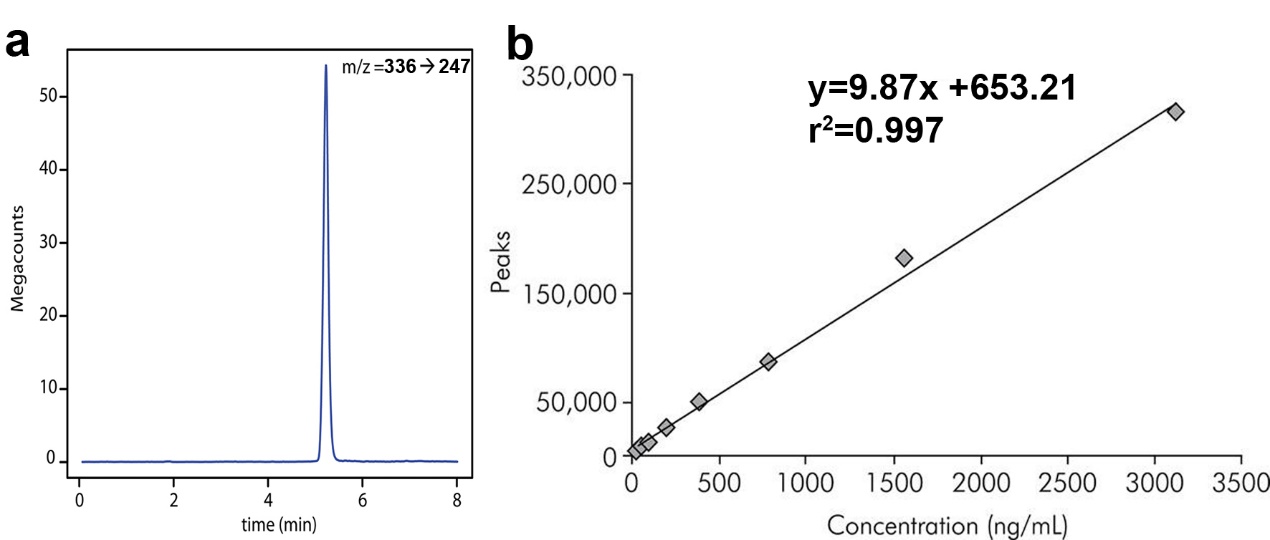

**Figure S6.** **(a)** Typical chromatograms based on the multiple reaction monitor (MRM) of hydroxychloroquine. **(b)** The standard curve for carboplatin. Calibration standards containing hydroxychloroquine in the concentrations (ng/mL) of 1, 5, 10, 200, 400, 800, 1600, and 3200 were prepared before injection into LC-MS/MS for analysis. Calibration curves of hydroxychloroquine exhibited excellent linearity over the concentration ranges of 1-3200 ng/mL.

**Evaluation of Drug loadings and their releasing profiles**

Hydroxychloroquine was quantified (for drug loading and releasing profile) using LC-MS/MS as described in *Supplementary information*. The drug loading (DL) of hydroxychloroquine in the freeze-dried nanoparticles powder was calculated using the following equation:

$$\boldsymbol{DL (\%)=}\frac{\boldsymbol{amount of hydroxychloroquine in freeze-dried nanoparticle (\mu g)}}{\boldsymbol{amount of freeze-dried nanoparticle (\mu g)}}\boldsymbol{\times100\%}$$

The entrapment efficiency (EE) was calculated from the following equation:

$$\boldsymbol{EE}\left( \boldsymbol{\%} \right)\boldsymbol{=}\frac{\boldsymbol{amount ofhydroxychloroquine in nanoparticle pellet}\left( \boldsymbol{\mu g} \right)}{\boldsymbol{amount of hydroxychloroquine in nanoparticle dispersion}\left( \boldsymbol{\mu g} \right)}$$

$$\boldsymbol{\times100\%}$$

Drug release study was carried out in physiological (PBS, pH 7.4) or acidic (acetate buffer, pH 5.5) and/or 20 mM dithiothreitol. 2 mL of 100 µg/mL of HCQ@ZnS@exo@iRGD were immersed in 100 mL of drug release media kept at 37 °C with constant stirring (80 rpm). Half milliliter of release media was taken at regular intervals and replaced with fresh media. After centrifugation, the amount of hydroxychloroquine in the release media was determined by LC-MS/MS.

**Cytotoxicity evaluation**

Cytotoxicity of different formulations was analyzed on U87 cells using MTT assay. Briefly, 1×10^4^ cells were seeded in 96 well plates for 24 h at 37°C and 5% CO_2_. After that, cells were incubated with free hydroxychloroquine or different hydroxychloroquine formulations at varying concentration range (6.25, 12.5, 25, 50, 100, and 200 µM) for 48 h with/without visible light irradiation. Control cells were maintained with media only. MTT solution (20 μL, 5 mg/mL) was added to each well and cultures were incubated for another 4 h. DMSO was added to dissolve the insoluble formazan crystal formed after MTT treatment, and absorbance was recorded at 550 nm using microplate reader (BioTek, Synergy H1 Hybrid reader). IC_50_ values were determined from these absorbance measurements.

**Caspase 3/7 activity assay**

Caspase 3/7 activity was performed using the Caspase-Glo 3/7 Assay (Promega) according to the manufacturer’s protocol. Cells were treated with different formulations and assayed after 24 h. Briefly, plates and reconstituted caspase 3/7 reagent were allowed to equilibrate to room temperature (RT), reagent was then added to wells at 1:1 ratio and incubated for 1 h at RT. Images of cells were taken by CLS. Wells with all assay components, except the cells, served as blank controls.

**Effect of HCQ@ZnS@exo@iRGD on 3D tumor spheroids**

BT-549 cells were allowed to form 3D spheroids by seeding them at a density of 5×10^3^ cells/100 μL per well in a Corning^®^ 96-Well Ultra-Low-Attachment Microplate and incubating them for several days. 3D spheroids were incubated for 24 h in culture medium containing free hydroxychloroquine, different hydroxychloroquine nanoparticles with/without light illumination or PBS (control). Then the supernatants were aspirated, fresh standard medium was added and cells were further cultured for another 5 days. On day 5, spheroids were photographed using phase-contrast microscopy.

**Autophagy inhibition decreases viability and stemness of U87 cells**

To verify the cytotoxic effects of HCQ@ZnS@exo@iRGD, the *in vitro* anti-tumor effects of free drugs and drug-loaded nanoparticles against U87 cells were tested using MTT assay. A significant increase in the percentage of dead U87 cells was observed when treated with HCQ@ZnS@exo (IC_50_=36.5±4.8 μm), compared with those treated with free HCQ (IC_50_=75.9±12.2 μm) or HCQ@ZnS (IC_50_=68.6±3.5 μm). Likewise, the number of viable cells decreased significantly upon the treatment with HCQ@ZnS@exo@iRGD (IC_50_=34.9±3.1 μm). Even greater cytotoxic effects were observed when U87 cells were treated with HCQ@ZnS@exo@iRGD under light illumination (IC_50_=12.6±1.9 μm) (Figure S7a). To visualize apoptotic events, cells were treated with the different formulations and then stained for apoptosis (cleaved caspase-3 (cCASP3)) markers. After the treatments, the percentage of cCASP3-positive U87 cells treated with HCQ@ZnS@exo@iRGD under light illumination significantly increased compared to the HCQ@ZnS@exo or non-illuminated HCQ@ZnS@exo@iRGD groups. In contrast, the apoptotic index of U87 cells in free HCQ and HCQ@ZnS were hardly affected by the treatments (Figure S7b).

Sphere formation assay, Boyden transwell chamber assay and would healing assay were performed to investigate the effects of the different formulations on the U87 self-renewal capacity and invasive activity, migratory activity. (Figure S7c, Figure S8a and b) The average number of spheroids formed under suspension cell culture conditions by HCQ@ZnS@exo and HCQ@ZnS@exo@iRGD treated U87 cells was significantly lower than the number formed by free HCQ and HCQ@ZnS. The sizes of the spheroids were also reduced. It was also noted that light irradiation further enhances the effect of the HCQ@ZnS@exo@iRGD in breaking up the spheroid. Using a Boyden transwell chamber assay, it was found that treating U87 cells with HCQ@ZnS@exo@iRGD under light illumination significantly reduced the percentage of those invasive cells by counting the numbers of cells on the lower surface of filter (Figure S8a). Such treatment also reduced the ability of the cells in a monolayer to fill the gap created by pipette scratching in a wound healing assay (Figure S8b).

These data show that U87 cell viability, stemness and invasive activity, migratory activity were strongly suppressed after HCQ@ZnS@exo@iRGD treatment under light irradiation.

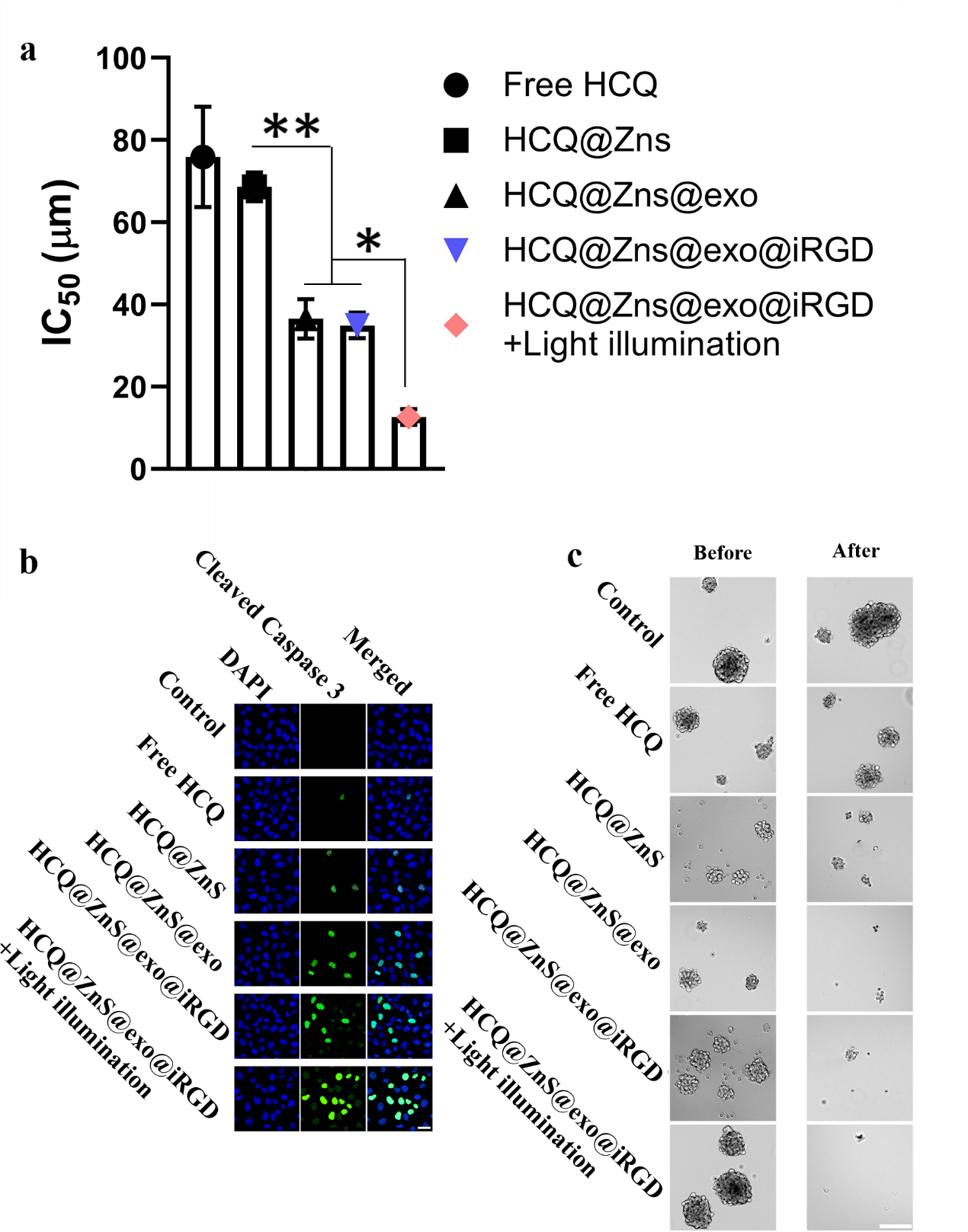

**Figure S7.** Cell viabilities, apoptosis and stemness of U87 cells after treatment with different formulations *in vitro*. **(a)** The IC_50_s for different formulations under different circumstance. Inhibiting autophagy reduces the viability of U87 cells. **(b)** Representative cleaved caspase 3 (green) immunofluorescence of U87 cells after treatment with the different formulations with/without light illumination. Scale bar is 20 μm. **(c)** Suspension culture conditions was used to assess the self-renewal capacity of U87 cells treated with different hydroxychloroquine formulations. The sphere number and cell number per sphere were counted after 7 days. Scale bar = 100 μm. All results are presented as data from three independent experiments. The data in panels are plotted as the mean ± SD with the indicated significance (*P < 0.05, **P < 0.01, according to a Student’s *t*-test).

**Effect of HCQ@ZnS@exo@iRGD on cell invasion and migration *in vitro***

In experiments to measure cell invasion, U87 cells (5 × 10^4^) in 300 µL serum-free DMEM were seeded into the upper chambers of 24-well cell culture plates with polycarbonate inserts with pore diameters of 8 µm (Millipore, Billerica, MA, USA). After a 24-h incubation, 800 µL of DMEM containing 10% serum was added to the lower chamber. The cells were then incubated for 24 h with PBS, free HCQ, HCQ@ZnS, HCQ@ZnS@exo or HCQ@ZnS@exo@iRGD, after which supernatants were aspirated, fresh standard medium was added, and cells were cultured for another 48 h. Cells that had not penetrated the filter were removed using cotton swabs, and cells that had penetrated to the lower surface of the filter were fixed with 4% paraformaldehyde and stained with 0.1% crystal violet. Cells were counted in five randomly selected fields at 40× magnification (Olympus, Japan). Each assay was performed in triplicate.

In experiments to measure cell migration, U87 cells were cultured to confluence for 24 h in 6-well plates in a 5 % CO_2_ incubator. The cell monolayer was scratched with a 200 µL pipette tip, then washed twice in PBS to remove floating cells. Cells were then cultured for 6 h in DMEM containing 0.5% FBS in the presence of PBS, free HCQ, HCQ@ZnS, HCQ@ZnS@exo or HCQ@ZnS@exo@iRGD. Supernatants were aspirated, fresh standard medium was added, and cells were cultured for another 18 h. Cells migrating from the leading edge were photographed under a light microscope.

Using a Boyden transwell chamber assay, we found that treating U87 cells with HCQ@ZnS@exo@iRGD under light illumination significantly attenuated the invasiveness of cells. (Figure S8a) In the wound healing assay, the gap change was little compared to the original width in cultures treated with HCQ@ZnS@exo@iRGD under light illumination. The gap was nearly completely covered in the control cells left untreated and in cells treated with free HCQ and HCQ@ZnS after 18 h of incubation. (Figure S8b)

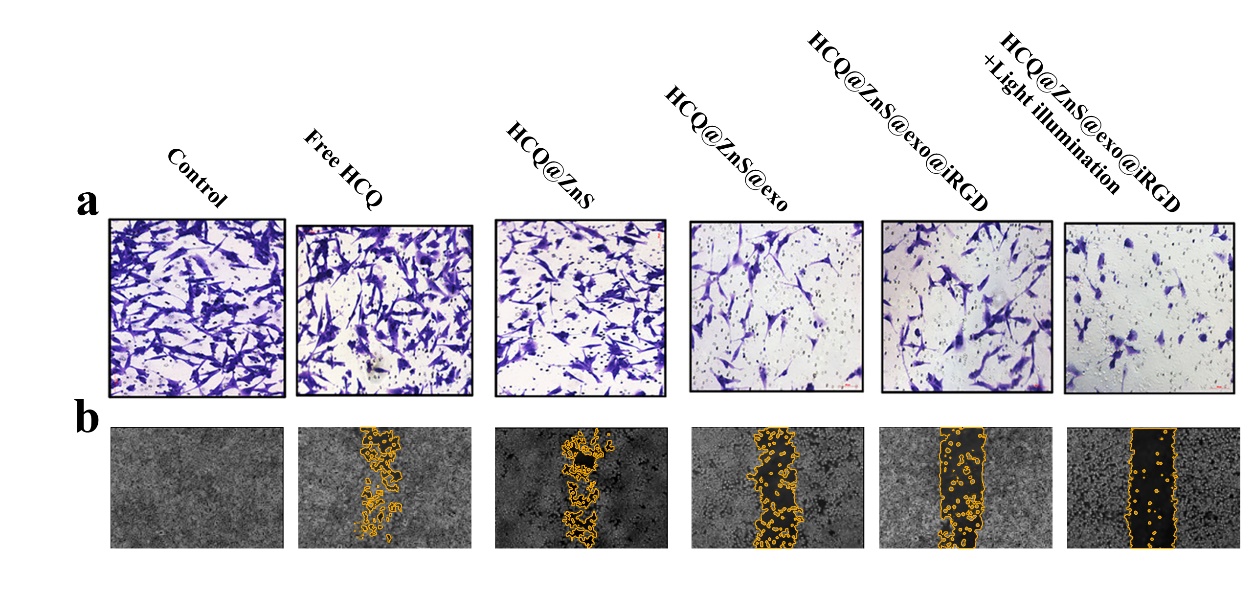

**Figure S8.** HCQ@ZnS@exo@iRGD effects on invasion and migration in U87 cells. **(a)** Typical images showing inhibition of U87 cell invasion by different formulations (magnification, 100×). **(b)** Cells were scratched with a pipette tip, then treated with free HCQ, HCQ@ZnS, HCQ@ZnS@exo or HCQ@ZnS@exo@iRGD with/without light illumination. The control group was left untreated. After incubation in fresh culture medium for another 18 h, migration was photographed under phase-contrast microscopy (magnification, 20×).

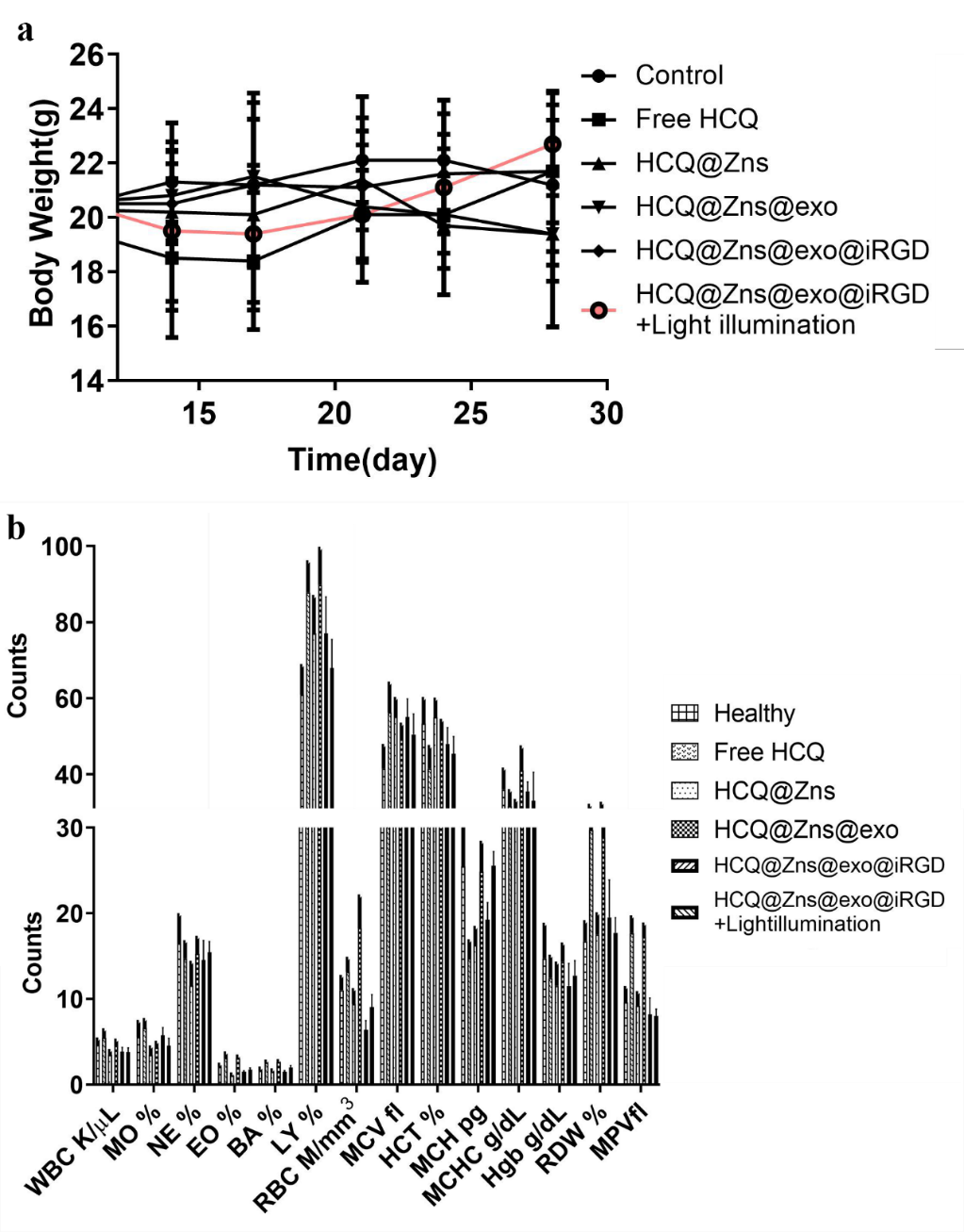
**Preliminary evaluation of side effects *in vivo***

**Figure S9.** Toxicity of different treatment groups showing no significant systemic damages. **(a)** Cumulative body weight during treatments. **(b)** Hematological changes in blood of different treatment groups. WBC, white blood cells; LY, lymphocytes; MO, monocytes; NE, neutrophils; EO, eosinophils; BA, basophils; RBC, red blood cells; MCV, mean corpuscular volume; HCT, hematocrit; MCH, mean corpuscular hemoglobin; MCHC, mean corpuscular hemoglobin concentration; Hgb, hemoglobin; RDW, red blood cell distribution width; MPV, mean platelet volume. Healthy group was treated with saline.
